# Molecular basis of tRNA modification by the human m^5^C methyltransferase NSUN2

**DOI:** 10.64898/2026.03.16.712100

**Authors:** Elodie Carmen Leroy, Michelangelo Lassandro, Arianna Di Fazio, Alessio Di Ianni, Kanhaya Lal, Jon Rodríguez-Villa, Alice Rossi, Andrea Graziadei, Monika Gullerova, Ana Casañal

**Author notes:** These authors contributed equally to this work (Elodie Carmen Leroy, Michelangelo Lassandro).

## Abstract

RNA 5-methylcytidine (m^5^C) is a prevalent modification that drives RNA stability and function. In humans, m^5^C is deposited on distinct RNA substrates by DNMT2/TRDMT1 and the NSUN family, to regulate diverse cellular processes, but how m^5^C writers recognise their substrates remains unclear. NSUN2 is a major m^5^C methyltransferase with broad roles in cell physiology and strong links to cancer and neurodevelopmental disorders ^1^. Here, we reconstitute an active human NSUN2-tRNA complex and capture its post-catalytic, tRNA-bound structure at 3.1 Å resolution. Using an integrated approach combining biochemistry, cryo-electron microscopy, crosslinking mass spectrometry and molecular dynamics simulations, we show that NSUN2 remodels the tRNA to access the variable-loop target cytidine. Recognition is driven by RNA architecture, with NSUN2 exploiting the L-shaped tRNA scaffold to position the target base in the catalytic centre. We further show that Gly679 at the NSUN2-tRNA interface is important for the stability of the complex, providing a mechanistic basis for how the disease-associated Gly679Arg substitution can impair tRNA binding. Together, these findings establish an RNA-structure-guided mechanism for NSUN2 substrate recognition and methylation and provide general principles for m^5^C deposition on cellular RNAs and their fundamental role in disease.

## MAIN

RNA modifications provide a rapid and reversible layer of gene expression regulation by modulating RNA structure and RNA-protein interactions. Their dysregulation leads to multiple human diseases ^2–5^. 5-methylcytidine (m^5^C) is widespread and conserved across eukaryotes and archaea, with established roles in RNA metabolism and translation ^1,6^, particularly in rRNAs and tRNAs, where it supports folding and stability ^6^.

In humans, m^5^C deposition is catalysed by DNMT2/TRDMT1 and the NOL1/NOP2/SUN (NSUN) family (NSUN1-7), a group of related m^5^C methyltransferases (‘writers’) that localise to distinct cellular compartments and thus act on nuclear/nucleolar, cytosolic and mitochondrial RNAs ^7,8^. These enzymes share a conserved Rossmann-fold catalytic core that binds S-adenosylmethionine (SAM) ^1^, yet structural views of their engagement with RNAs remain limited. Structural snapshots exist for NSUN1 and NSUN4 in ribosome and mito-ribosome assembly intermediates, and for NSUN6 in complex with tRNA and mRNA, but how substrate specificity is achieved across the NSUN-family remains unknown ^9–18^.

NSUN2 is a major, clinically relevant m^5^C writer that regulates proliferation, differentiation and stress responses ^8,19^. Loss of NSUN2-dependent tRNA methylation promotes stress-induced tRNA cleavage and reduces translation ^20^. Furthermore, NSUN2 loss-of-function variants cause neurodevelopmental disorders, including autosomal-recessive intellectual disability and Dubowitz syndrome ^7,21,22^. NSUN2 modifies many cytosolic tRNAs and additional substrates such as mRNAs and vault RNA ^23–26^. In tRNAs, NSUN2 predominantly targets variable loop cytidines (C48/49/50), complementing DNMT2 (C38) and NSUN6 (C72) ^27,28^. Here, we provide structural and functional insight into NSUN2-tRNA recognition, clarifying how a broad-spectrum m^5^C writer accesses variable-loop cytidines within an intact tRNA scaffold, and offering a structural rationale for a disease-associated variant at the tRNA-binding interface.

## RESULTS

### Post-catalytic cryo-EM structure of the human NSUN2-tRNA^Asp^_GUC_ complex

To define how human NSUN2 positions variable-loop cytidines on tRNA for methyl transfer, we reconstituted an active NSUN2-tRNA complex and determined its structure by cryo-electron microscopy (cryo-EM) at 3.1 Å resolution (Fig. 1, Extended Data Fig. 1). To stabilise a post-catalytic intermediate ^29,30^, we used an NSUN2 Cys271Ser mutant (C271S). This construct exploits the two-cysteine turnover mechanism ^17,31^, in which Cys321 forms the covalent enzyme-RNA intermediate and Cys271 promotes its resolution (Fig. 1a). The structure was solved in complex with human tRNA^Asp^_GUC_, a well-characterised NSUN2 substrate carrying variable-loop cytidines C47 and C48 (Fig. 1b) ^20,32^. Electrophoretic mobility shift assays (EMSAs) showed that wild-type NSUN2 and NSUN2^C271S^ bound tRNA^Asp^_GUC_ with comparable micromolar affinity (Extended Data Fig. 2). Incubation of NSUN2^C271S^ with tRNA^Asp^_GUC_ and SAM yielded a covalent complex that persisted under denaturing SDS-PAGE conditions and had an apparent mass of ∼115 kDa by mass photometry, consistent with a 1:1 NSUN2-tRNA complex ratio (Fig. 1c).

**Figure 1.**
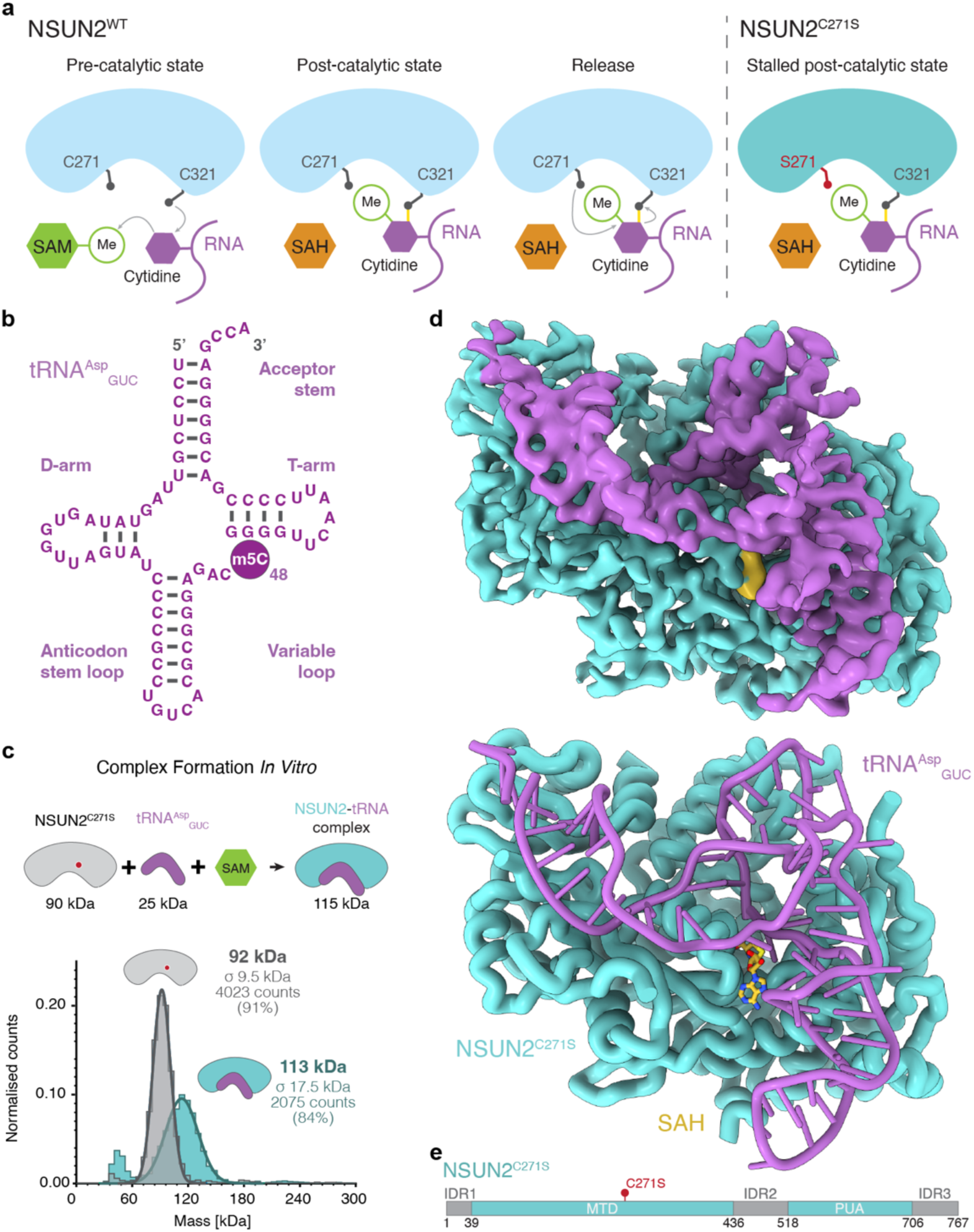
Structure of human NSUN2-tRNA^Asp^_GUC_ in a post-catalytic state. **a,** Schematic of the NSUN2 catalytic cycle highlighting the two-cysteine turnover mechanism, in which Cys321 forms a covalent NSUN2–RNA intermediate and Cys271 promotes its resolution; substitution Cys271Ser stalls the enzyme in a post-catalytic state after methyl transfer. **b,** Secondary structure of human tRNA^Asp^_GUC_ indicating the variable-loop target cytidine and the methylation site at C48 (m^5^C48). **c,** *In vitro* reconstitution of the covalent NSUN2^C271S^-tRNA^Asp^_GUC_ complex in the presence of S-adenosylmethionine (SAM) and mass photometry analysis showing monomeric NSUN2 and a higher-mass species consistent with a 1:1 NSUN2–tRNA complex. **d,** Cryo-EM reconstruction and model of the NSUN2^C271S^-tRNA^Asp^_GUC_ complex, in which the entire tRNA is visualised and runs along one face of NSUN2, engaging an extended surface formed by the MTD and PUA domains (NSUN2 in teal; tRNA in purple; SAH in yellow). **e,** Domain architecture of human NSUN2 with the methyltransferase domain (MTD) and PUA RNA-binding domain indicated; intrinsically disordered regions (IDR1-IDR3) are shown as not resolved in the reconstruction.

To obtain a high-resolution reconstruction of the NSUN2-tRNA^Asp^_GUC_ complex, we collected cryo-EM datasets from non-crosslinked and mildly crosslinked samples prepared using sulfosuccinimidyl 4,4′-azipentanoate (sulfo-SDA; sSDA) within our photo-crosslinking cryo-EM workflow (pXL-EM, also used in a related preprint ^33^), which stabilises the complex prior to grid preparation and enhances map quality (Fig. 1d, Extended Data Fig. 1). The non-crosslinked dataset adopted the same overall architecture but was limited by conformational flexibility, and we therefore focused subsequent processing on the sSDA-treated sample. Both NSUN2-tRNA preparations (with and without crosslinking) contained a mixture of monomeric NSUN2 and higher-molecular mass oligomeric species (including dimers), consistent with the ability of NSUN2 to form multiple assembly states (the purification and reconstitution workflow is described in Extended Data Fig. 2).

The cryo-EM reconstruction resolves the structure of human NSUN2 bound to tRNA^Asp^_GUC_ in a 1:1 complex (Fig. 1d). NSUN2 comprises a Rossmann-like SAM-dependent methyltransferase domain (MTD) and a PUA RNA-binding domain (Fig. 1e). Three regions of NSUN2 predicted to be intrinsically disordered (IDR), located at the N-terminus, C-terminus and in the linker between the MTD and PUA domains, are not resolved in our cryoEM map, consistent with AlphaFold models (Fig. 1e; Supplementary Data Fig. 1).

The MTD and PUA domains together form an extended RNA-binding surface that engages the tRNA along its length: the MTD binds the T-arm, variable loop and anticodon stem-loop, whereas the PUA domain binds the acceptor stem, together spanning the full length of the tRNA (Fig. 1d). In the catalytic centre, the target cytidine C48 is covalently linked to Cys321, and clear density is present for S-adenosylhomocysteine (SAH), confirming capture of a post-methyl-transfer state (Extended Data Fig. 3).

To aid model building in locally lower-resolution and flexible regions, we integrated complementary crosslinking mass spectrometry (MS) datasets. Photo-crosslinking and sulfo-SDA provided protein-protein restraints that corroborated the overall architecture of the complex and supported modelling of the PUA domain as a flexible platform for the tRNA acceptor stem (Extended Data Fig. 1,4). UV RNA-protein crosslinking supported confident placement of the tRNA and revealed the pronounced flexibility of the anticodon stem (see Methods; Extended Data Fig. 5, Supplementary Fig. 4).

Together, these data define the architecture of the NSUN2^C271S^-tRNA^Asp^_GUC_ complex and provide a structural basis for understanding how NSUN2 tRNA recognition and methylation.

### NSUN2 remodels tRNA^Asp^_GUC_ to position the variable loop for catalysis

A mature tRNA adopts the canonical L-shaped architecture in which the variable loop forms stabilising long-range tertiary contacts with the hinge/D-arm junction and the D-arm–acceptor stem, together defining the characteristic tRNA elbow fold (Fig. 2 a,b) ^34^. Interestingly, in the NSUN2-bound state, tRNA^Asp^_GUC_ adopts a substantially remodelled, more open conformation that matches the enzyme surface (Fig. 2c). The native tertiary interactions that tether the variable loop to the D-loop and hinge are relaxed, and the D-loop itself loses its ordered conformation. The hinge disengages from the D-arm and becomes uncoupled from the tRNA core, consistent with reduced local cryo-EM density in this region (Extended Data Fig. 1). By contrast, the D-arm remains folded and continues to associate with the T-arm, and the acceptor stem and anticodon stem-loop retain near-canonical conformations.

**Figure 2.**
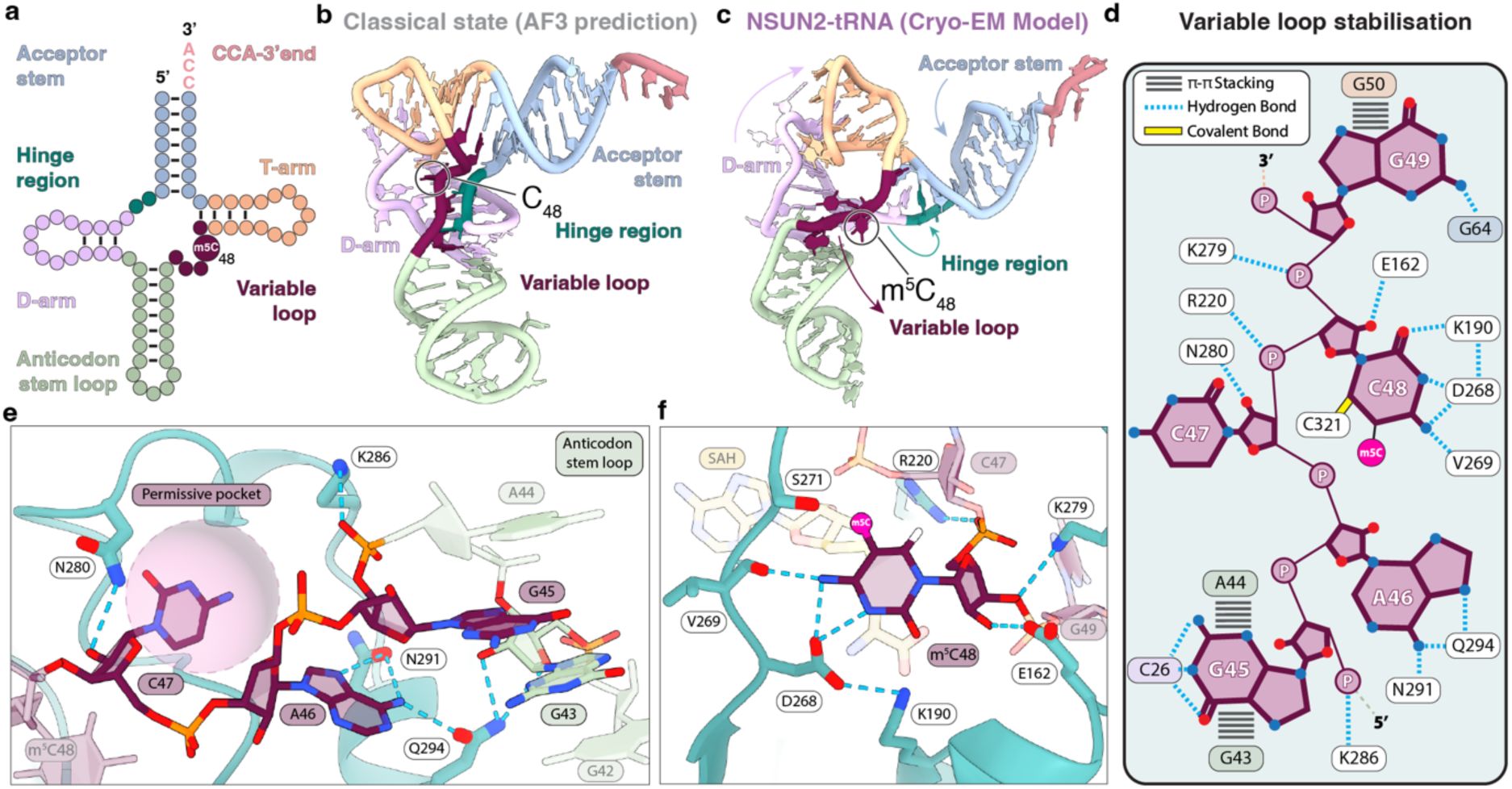
tRNA remodelling for methylation. **a,** Clover-leaf secondary structure representation of tRNA^Asp^_GUC_ elements highlighting acceptor arm (light blue), hinge region (green), D-arm (pink), anticodon arm (light green), variable loop (magenta), T-arm (light orange), and CCA-3′ end (salmon). **b,** Classical tRNA conformation (AlphaFold3 prediction) showing the position of C48 within the canonical L-shaped fold. **c,** Cryo-EM model of the NSUN2-bound tRNA^Asp^_GUC_ showing a remodelled, more open tRNA conformation in which the variable loop is redirected toward the NSUN2 active site. **d,** Interaction schematic summarizing variable-loop contacts and active-site engagement; m^5^C48 is highlighted in fuchsia, and the covalent linkage between C48 and Cys321 is highlighted in yellow. **e,** Close-up of the upstream variable-loop region highlighting a permissive binding pocket accommodating the nucleotide at position 47 and stabilizing interactions that guide the loop toward the catalytic centre. **f,** Close-up of the post-catalytic active site showing positioning of m^5^C48 (covalently trapped in the NSUN2^C271S^ complex) and surrounding catalytic and RNA-stabilizing residues, with SAH bound in the cofactor pocket.

Collectively, these rearrangements release the variable loop from the tRNA core. The loop is no longer embedded within the tRNA elbow but instead projects toward the catalytic pocket, with nucleotides 44-48 adopting an extended geometry that enables insertion into the active site (Fig. 2d). Accordingly, the NSUN2 catalytic centre accommodates cytidines presented within the short variable loop characteristic of type I tRNAs. At the 5′ end of the variable loop, A46 is withdrawn from its interaction with the G22-U13 wobble pair and reoriented by Gln294 and Asn291 to open the loop upstream of the methylation site, while Lys286 clamps the G45 phosphate to anchor the upstream segment on the MTD surface (Fig. 2e). This remodelling disrupts the conserved Levitt base pair (a long-range interaction that normally tethers C47 to G15) and replaces it with an alternative anchor, as Asn280 engages the C47 O2′ to stabilise the loop at the pocket entrance and guide its trajectory toward the active site (Fig. 2e). The pocket accommodating C47 is permissive to alternative nucleobases, supporting a dominant role for RNA architecture over sequence identity at position 47 (Extended Data Fig. 6).

Within the catalytic centre, C48 unpairs from G63 in the T-arm and is positioned for methyl transfer (Fig. 2f). In the trapped post-catalytic complex, m^5^C48 is covalently linked to Cys321 and inserted into an active-site pocket formed by Cys321, Lys190, Asp268 and Ser271 (C271S), adjacent to the cofactor-binding site (Fig. 2f). Asp268 and Val269 contact the cytidine base, providing a structural rationale for cytidine selectivity at the target position (Fig. 2f). The catalytic centre geometry is conserved relative to NSUN6 (PDB: 5WWS ^17^, Extended Data Fig. 7), supporting a shared catalytic strategy across NSUN enzymes.

### NSUN2 anchors the conserved tRNA elbow while accommodating local remodelling

Despite extensive remodelling to liberate the variable loop, NSUN2 recognises conserved features shared among canonical tRNAs (Fig. 3). The overall T-arm architecture is preserved and the D-T contacts that maintain the L-shaped fold remain largely intact (Fig. 3a,b).

**Figure 3.**
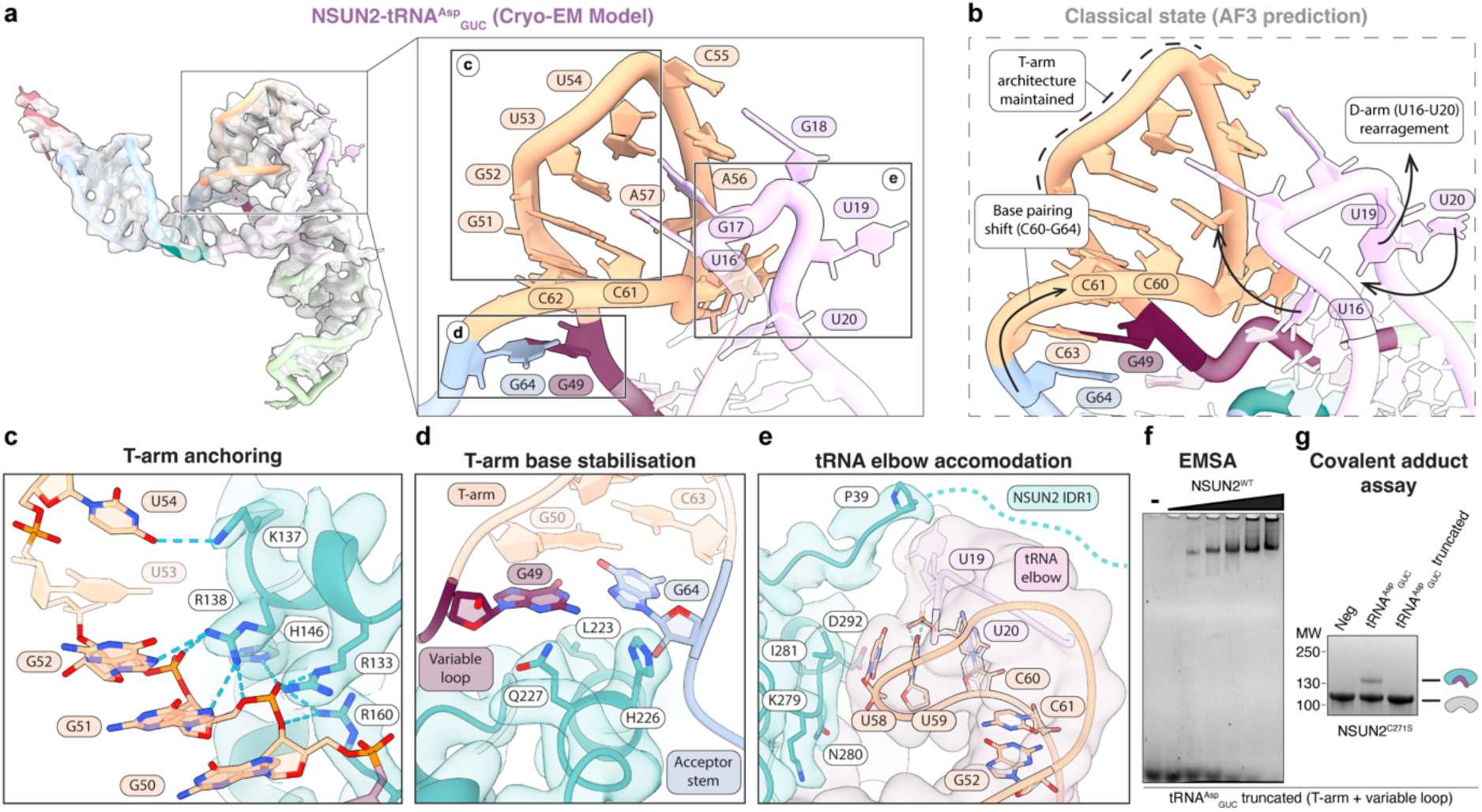
Elbow anchoring with local plasticity. **a,** Cryo-EM density with the fitted tRNA^Asp^_GUC_ model, with close-up views of the tRNA elbow in the NSUN2-tRNA^Asp^_GUC_ structure. **b,** Comparison to a classical tRNA conformation (AlphaFold3 prediction) indicating preservation of overall T-arm architecture alongside rearrangements in the D-arm/hinge region and a local base-pairing register shift in the T-arm stem (C60–G64 region). **c,** NSUN2 T-arm anchoring interface showing electrostatic and hydrogen-bonding interactions that stabilize nucleotides around G50–G52 and a base-specific contact to the conserved U54. **d,** Stabilization of the T-arm immediately downstream of the variable loop, showing a protein platform that supports a local register shift while maintaining stem integrity. **e,** Accommodation of the remodelled elbow, including repositioning of the U58-U59-C60 loop. **f,** Electrophoretic mobility shift assay (EMSA) assessing binding of NSUN2 to a truncated tRNA comprising the T-arm fused to the variable loop showing that RNA binding is maintained (see Extended Data Fig. 1 for EMSA assays of NSUN2 with full-length tRNA^Asp^_GUC_). **g,** Covalent adduct formation assay showing that the truncated T-arm+variable-loop substrate does not support productive covalent complex formation under the conditions tested, in contrast to full-length tRNA^Asp^_GUC_.

NSUN2 recognises the T-arm immediately downstream of the variable loop through a cluster of positively charged residues that surrounds the phosphate backbone and forms electrostatic and hydrogen-bonding contacts around G51-G52, stabilising nucleotides 49-53 (Fig. 3c). In addition, Lys138 makes a base-specific contact to U54, a highly conserved nucleotide typically modified to ribothymidine, highlighting a conserved feature characteristic of mature tRNAs at the elbow (Fig. 3c).

NSUN2 also stabilises local plasticity within the anchored elbow: opening of the D-arm-acceptor-stem junction extends the tRNA and shifts the elbow onto the NSUN2 surface to accommodate the binding site (Fig. 3b, d, e). Immediately downstream of the target site (C48) Leu 223, His 226, and Gln227 form a platform contacting G49 and G64, supporting a local register shift in which G49 pairs with G64 instead of the canonical G49-G63 pairing, allowing C48 to be released for catalysis while maintaining T-arm integrity (Fig. 3d). This re-pairing occurs within a GC-rich segment, buffering elbow rearrangements without disrupting the global T-arm scaffold.

At the top of the elbow, the U58-U59-C60 loop shifts towards the D-arm, C60 flips out and redirects toward U20, forming stacking and hydrogen-bonding contacts (Fig. 3e). This rearranged loop packs against the NSUN2 surface near residues 279-292, stabilising the remodelled elbow and preserving the overall D-T tertiary core and L-shaped geometry (Fig. 3e). UV RNA-protein crosslinking supports the assignment of an extra density near the elbow as the N-terminal region of NSUN2 (IDR1), (Extended Data Fig. 5, Supplementary Data Fig. 4), suggesting that flexible elements further reinforce this anchored configuration.

To test whether the T-arm anchoring module is sufficient for catalysis, we engineered a minimal RNA comprising the tRNA^Asp^_GUC_ T-arm fused to the variable loop. NSUN2 bound this truncated substrate in EMSAs (Fig. 3f), but it did not support covalent adduct formation in catalytic assay (Fig. 3g), indicating that T-arm binding alone is insufficient for catalysis. These results support a model in which NSUN2 uses conserved elbow/T-arm features as an anchoring platform while tolerating local remodelling but requires additional contacts from an intact tRNA fold for productive catalysis.

### NSUN2 is structurally adapted to engage tRNA-like architectures

To define the additional contacts required beyond T-arm anchoring, we mapped how NSUN2 engages the full-length tRNA through distributed contacts (Fig. 4).

**Figure 4.**
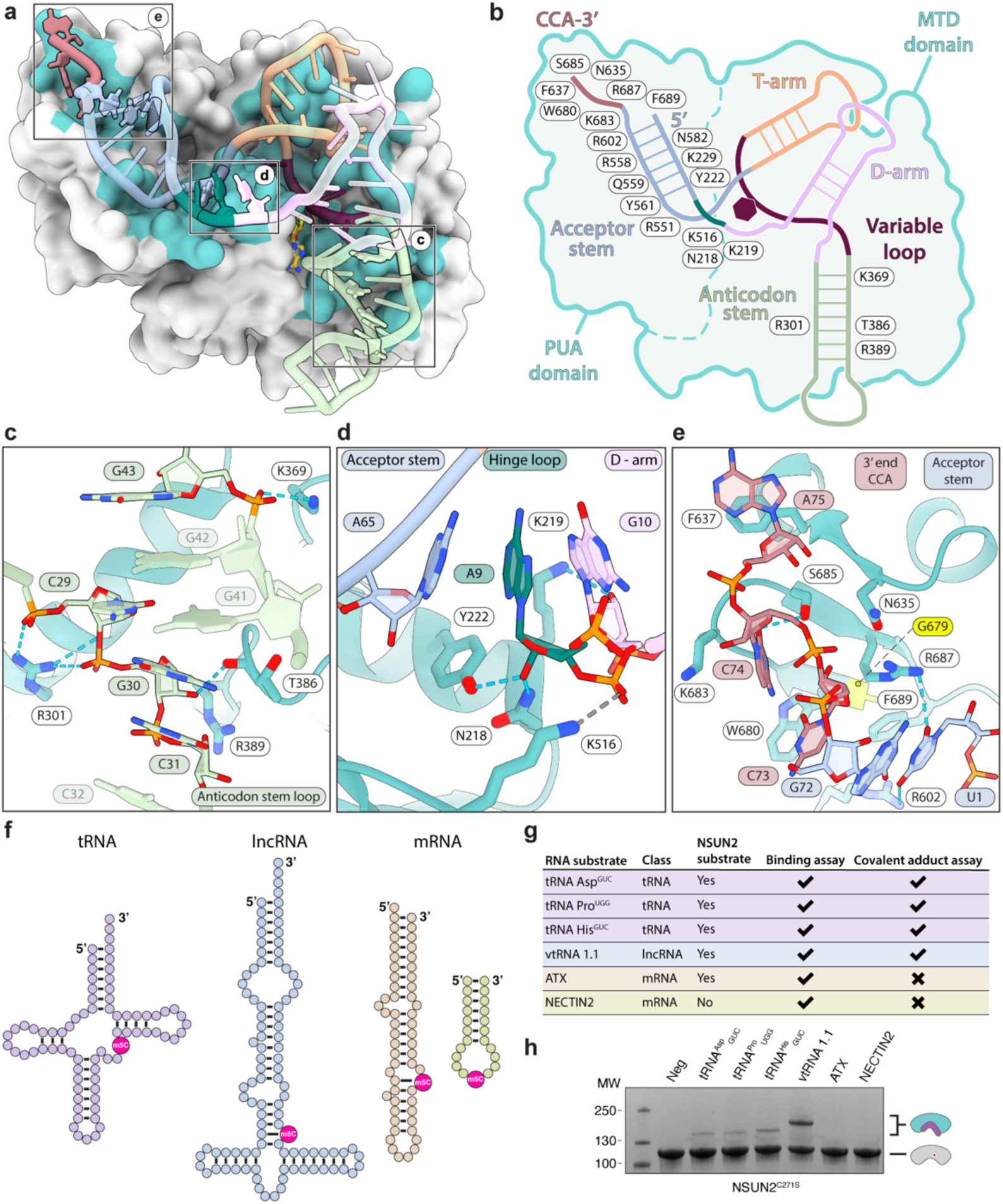
tRNA-like fold recognition by NSUN2. **a,** Overview of NSUN2 showing an extended RNA-binding surface spanning the methyltransferase domain (MTD) and PUA domain, with residues contributing to this interface highlighted in blue. **b,** Schematic of human NSUN2 bound to tRNA^Asp^_GUC_ showing multi-point engagement of the tRNA with the MTD and PUA domains of NSUN2. **c,** Close-up of anticodon stem-loop engagement by the MTD, dominated by backbone-mediated interactions. **d,** NSUN2 clamps the hinge-region backbone to stabilise the opened junction. **e,** The PUA domain recognises the mature tRNA termini, stabilising the 5′ end (U1) and anchoring the 3′ CCA tail through stacking and hydrogen-bonding interactions; the disease-associated variant Gly679Arg (G679R) lies adjacent to this site and is predicted to perturb local packing and weaken terminus recognition. **f-h,** RNA-binding and activity profiling across diverse RNA substrates: electrophoretic mobility shift assays show that NSUN2 associates broadly with RNA, whereas covalent-adduct formation and methylation are robust primarily on canonical tRNAs and vault RNA 1.1, consistent with a requirement for a tRNA-like fold that correctly presents the variable loop to the catalytic centre.

NSUN2 presents an extended, positively charged RNA-binding surface spanning the MTD and PUA domains, enabling multi-point engagement with the entire tRNA structure (Fig. 4a,b). The NSUN2-tRNA^Asp^ interface buries ∼2690 Å² and is supported by extensive hydrogen-bonding interactions (Supplementary Table S4).

Within the MTD, NSUN2 anchors the anticodon stem-loop primarily through backbone contacts: Arg301 engages the phosphates of C29 and G30 (and the O3’ of C29), while Thr386 and Lys369 contact the backbone of nucleotides C31 and G43 (Fig. 4c). At the acceptor-stem-D-arm junction (hinge region), NSUN2 provides an additional anchoring point, with Lys219 and Lys516 clamping the phosphate backbone of nucleotides 9-11, stabilising the opened junction through a stacking bridge that links the acceptor stem (A65), the hinge loop (A9) and the D-arm (G10) (Fig. 4d). This composite stacking network, together with phosphate clamping, locks the extended hinge conformation. Alongside MTD-mediated sequestration of the variable loop and elbow region, it stabilises the tRNA geometry that presents the target cytidine to the catalytic centre.

NSUN2 also forms a defined binding pocket for the mature tRNA termini: at the 5’end, U1 is stabilised by stacking against Phe689 and hydrogen bonding with Arg602/Arg687, supporting the U1-G72 wobble pair (Fig. 4e). At the 3’end, the CCA tail is anchored by stacking and polar contacts (Trp680-C73, Asn635, Ser685 Lys683-C74, and Phe637-A75; Fig. 4e), highlighting the capacity of NSUN2 to engage CCA-containing tRNAs late in the maturation pathway. The PUA domain also tracks along the acceptor stem via backbone contacts on the 3′ strand, further stabilising the acceptor-stem/CCA module (Fig. 4b).

Notably, the Dubowitz syndrome-associated NSUN2 missense variant Gly679Arg (G679R), ^21,22,35^ maps to a β-strand in the PUA domain immediately adjacent to the site that accommodates the mature tRNA ends (Fig. 4e). In the structure, this region lies next to the CCA-tail interface and the 5′-end pocket that supports the U1-G72 wobble pair and the U1-Phe689 stacking interaction (Extended Data Fig. 8a,b). We therefore hypothesised that G679R perturbs the geometry of this termini-binding pocket (Extended Data Fig. 8c,d). To assess the impact of the substitution, we performed molecular dynamics (MD) simulations of the NSUN2^C271S^-tRNA^Asp^_GUC_ complex, initiated from the cryo-EM model, and introduced G679R. MD simulations (up to 500 ns) revealed a significant reduction in complex stability in the presence of G679R. This was driven by displacement of a loop near the termini pocket, which disrupts the U1-Phe689 stacking interaction observed in the wild-type structure (Extended Data Fig. 8e,f). These results indicate that G679R compromises productive NSUN2-tRNA binding, in agreement with the reduced tRNA binding and activity reported for this variant ^36^.

Finally, we compared RNA binding with catalysis. NSUN2 associated with diverse RNAs *in vitro* (Fig. 4f, Extended Data Fig. 9), yet productive covalent-adduct formation required an intact variable-loop methylation site: a tRNA^Gly^_CCC_ mutant in which variable loop cytidines C46-48 were substituted with A46-48 bound NSUN2 comparably to wild-type tRNA^Gly^_CCC_, but failed to form a covalent complex under trapping conditions (Extended Data Fig. 9). Consistently, covalent-adduct formation and methylation were robust primarily on canonical tRNAs and vault RNA 1.1, whereas short mRNA fragments remained unmethylated (Fig. 4g,h). These results support a model in which NSUN2 can bind RNAs promiscuously, but methyl transfer requires multi-point recognition of a tRNA-like structure together with a correctly configured variable-loop cytidine motif.

## DISCUSSION

The NSUN2-tRNA structure provides a direct view of how a broadly RNA-binding m^5^C writer achieves selective chemistry: RNA association is promiscuous, whereas productive methyl transfer requires multi-point recognition of a tRNA-like architecture. Consistently, NSUN2 binds diverse RNAs *in vitro*, yet robust covalent-adduct formation and methylation are largely restricted to structured substrates such as canonical tRNAs and vault RNA 1.1, indicating that binding alone is insufficient for turnover.

A central mechanistic feature of the complex is the tRNA remodelling required to access the variable loop. Rather than engaging a pre-formed elbow, NSUN2 exploits tRNA conformational plasticity to loosen elbow packing and deliver a short variable-loop segment to the active site while preserving the L-shaped scaffold. This tRNA remodelling, key to the recognition of tRNA, was missed in predictions by AlphaFold3. Although remodelling is common in tRNA-processing and modification enzymes ^34^, NSUN2 stabilises an unusually remodelled state by clamping conserved elbow/T-arm landmarks and using distributed contacts, enabling broad tRNA recognition without sacrificing catalytic stringency.

At the level of chemistry, NSUN2 uses the conserved SAM-dependent Rossmann-fold architecture shared by many RNA methyltransferases, including METTL1-WDR4 (m^7^G) and the METTL3-METTL14 complex (m^6^A) ^37,38^. What distinguishes NSUN2 is therefore substrate presentation: a broad, distributed MTD-PUA interface supports RNA engagement and productive alignment of the target cytidine. NSUN6 also adopts an MTD-PUA organisation, but its domains are more compact and provide a smaller RNA-contact surface ^16,17^, whereas METTL1-WDR4 relies on an accessory factor and a distinct tRNA interactions ^37,39^.

Our structure also clarifies how disease-associated substitutions compromise NSUN2 function. The Dubowitz syndrome-linked G679R substitution maps adjacent to the peripheral surface that helps secure the mature tRNA termini. Consistent with this placement, MD simulations indicate that G679R destabilises the NSUN2-tRNA assembly and increases conformational lability. This supports a model in which G679R impairs the architecture required for stable, catalytically competent substrate engagement. More broadly, disruption of peripheral recognition and anchoring surfaces, rather than the conserved catalytic core, can compromise the architectural prerequisites for catalysis and contribute to disease. Finally, our structure and simulations provide a foundation for structure-guided NSUN2 inhibitor development, expanding strategies beyond the SAM-binding pocket to peripheral RNA-contact and tRNA-anchoring surfaces ^40,41^.

In addition to the monomeric 1:1 NSUN2-tRNA complex resolved here, NSUN2 can form higher-molecular mass species *in vitro*, but heterogeneity in 2D classes and low-resolution reconstructions precludes mechanistic interpretation (Extended Data Fig. 2). Previous work has shown that glucose binding can enhance NSUN2 oligomerisation and promote NSUN2-m^5^C deposition and associated phenotypes in cancer and epidermal differentiation contexts ^42,43^. If regulated in cells, oligomerisation could tune localisation, substrate selection, or catalytic efficiency and will require dedicated structural and cellular analysis.

In summary, the NSUN2-tRNA structure defines an architecture-driven mechanism in which multi-point contacts remodel the tRNA elbow to bring the variable loop to the catalytic centre, providing a structural basis for NSUN2 substrate scope and for understanding how mutations at RNA-binding surfaces impair m^5^C deposition in human disease.

## Materials and Methods

### Expression and Purification of NSUN2^WT^

6xHis-NSUN2^WT^ was expressed using the Bac-to-Bac expression system (Thermo Fisher Scientific). 1L of low-passage (<30) Sf9 cells grown in suspension at 27 °C, 110 rpm in Sf-900 II SFM buffer (Gibco, 10902088) were infected with 5 mL of V_1_baculovirus stock. Cells were harvested 48 hours post-infection by centrifugation at 700 g for 15 minutes at 4 °C.

6xHis-NSUN2^WT^ was purified as follows, all steps were carried out at 4 °C. Cell pellets were lysed with 6 volumes of lysis buffer (50 mM HEPES pH 7.6, 500 mM NaCl, 10% glycerol, 0.5% Triton-X 100, 1 mM TCEP, 1x Complete protease inhibitor EDTA-free, 1x PMSF, and 1x leupeptin) and 2500 U of universal nuclease (Pierce), with rotation for 30 min. The lysate was then sonicated five times, 30 s on/off at 15 μm, spun at 13,000 *g* for 45 min, and the supernatant filtered with a 0.2-μm filter (Corning, 430049). Cleared lysate was then loaded onto Ni-NTA agarose resin (QIAGEN, 30210) in a gravity flow column pre-equilibrated with 5 volumes of wash buffer (50 mM HEPES pH 7.6, 500 mM NaCl, 5% glycerol, 1 mM TCEP, 1x Complete protease inhibitor EDTA-free, 1x PMSF, 1x leupeptin, and 10 mM imidazole) and washed with 20 volumes of wash buffer. Proteins were eluted with elution buffer (50 mM HEPES pH 7.6, 500 mM NaCl, 5% glycerol, 1 mM TCEP, 1x Complete protease inhibitor EDTA-free, 1x PMSF, 1x leupeptin, and 500 mM imidazole). Fractions containing NSUN2 were dialyzed overnight against Heparin low salt buffer (50 mM HEPES pH 7.0, 200 mM NaCl, 1 mM TCEP, 5 mM MgCl_2_, 5% glycerol) and then applied to a 5 mL HiTrap Heparin HP column (Cytiva) equilibrated in Heparin low-salt buffer. After washing with 20 CV of low-salt buffer, proteins were eluted using a linear gradient (0–65%) into high-salt buffer (50 mM HEPES pH 7.0, 1 M NaCl, 1 mM TCEP, 5 mM MgCl2, 5% glycerol), followed by a final step at 100% high-salt buffer. Purified fractions were pooled, concentrated, snap-frozen in liquid nitrogen, and stored at −80°C.

### Cloning and mutagenesis of the NSUN2^C271S^ construct

The coding sequence of human NSUN2 (UniProt accession Q08J23-1) was gene-synthesized and cloned by Epoch Life Science into the pACEBac1 vector, encoding a C-terminal Twin-Strep tag preceded by an HRV 3C protease cleavage site.

The NSUN2^C271S^ mutant construct was generated using the QuickChange II Site-Directed Mutagenesis Kit (Agilent Technologies) following the manufacturer’s instructions. Briefly, PCR reactions were performed with specific complementary primers (Supplementary Table S2) using a vector containing the NSUN2 wild-type gene tagged with TwinStrep at the C-terminus. PCR products were digested with DpnI (10 U/µL, NEB) for 1 h at 37 °C, then transformed into XL1-Blue competent cells. Transformants were selected on LB-agar plates supplemented with 50 µg/mL kanamycin. Plasmid DNA was isolated using the QIAprep Spin Miniprep Kit (Qiagen) and verified by whole-plasmid sequencing (Eurofins Genomics).

### Expression and Purification of NSUN2^C271S^

Recombinant NSUN2^C271S^-TwinStrep protein was produced using the MultiBac baculovirus/insect cell expression system. Recombinant bacmids were generated in EmBacY cells as previously described ^44^. Baculovirus stocks were amplified through two passages (P1 and P2) in Sf9 cells. For P1 production, Sf9 cells were seeded at 0.5 × 10⁶ cells ml⁻¹ in 2 ml per well of a six-well plate and transfected with bacmid DNA using Cellfectin II (Gibco) according to the manufacturer’s instructions. P1 virus was harvested 72 h post-transfection and supplemented with 40 μl newborn calf serum (Gibco) prior to storage at 4 °C.

For P2 amplification, Sf9 cells at 0.5 × 10⁶ cells ml⁻¹ were infected with the entire P1 vial in a final volume of 25 ml. The P2 virus was harvested 48 h post-infection and stored at 4 °C until use for protein expression.

For protein expression, High Five cells were grown in 500 mL Insect-XPRESS™ medium (Lonza Biosciences) containing 1% penicillin/streptomycin and infected with 2 ml of the corresponding baculovirus. Cultures were incubated for 48 h at 27 °C with shaking at 130 rpm. Cells were harvested by centrifugation at 4,000 × g for 10 min at 4 °C, and pellets were stored at −80 °C.

For protein purification, cell pellets were resuspended in lysis buffer containing 50 mM HEPES (pH 7.5), 200 mM NaCl, 1 mM TCEP, 1 mM EDTA, 5% (v/v) glycerol, cOmplete™ EDTA-free protease inhibitor (Merck), and benzonase. Cells were lysed by mechanical homogenization using a 100 mL Kimble tissue grinder. Lysates were clarified by centrifugation at 25,000 × g for 30 min at 4 °C. Clarified lysate was loaded onto a 5 mL StrepTrap XT column (Cytiva) pre-equilibrated in wash buffer (50 mM HEPES pH 7.5, 200 mM NaCl, 1 mM TCEP, 1 mM EDTA, 5% glycerol) using an ÄKTA pure system (Cytiva). After washing with 20 column volumes (CV), proteins were eluted with 8 CV elution buffer containing 50 mM biotin.

Eluted fractions were applied to a 5 mL HiTrap Heparin HP column (Cytiva) equilibrated in low-salt buffer (50 mM HEPES pH 7.5, 100 mM NaCl, 1 mM TCEP, 5 mM MgCl2, 5% glycerol). After washing with 20 CV of low-salt buffer, proteins were eluted using a linear gradient (0–65%) into high-salt buffer (50 mM HEPES pH 7.5, 1 M NaCl, 1 mM TCEP, 5 mM MgCl2, 5% glycerol), followed by a final step at 100% high-salt buffer. Peak fractions were concentrated and further purified by size-exclusion chromatography (SEC) on a Superdex 200 Increase 10/300 column (Cytiva) equilibrated in SEC buffer (50 mM HEPES pH 7.0, 150 mM NaCl, 1 mM TCEP, 5 mM MgCl₂). Protein concentrations were measured using a NanoDrop™ OneC spectrophotometer (Thermo Fisher Scientific). Final samples were flash-frozen in liquid nitrogen and stored at −80 °C. Purification steps were monitored by SDS–PAGE and Instant Blue Coomassie (Abcam) staining.

### Mass photometry

Mass photometry measurements were performed using a Refeyn TwoMP instrument. Proteins were diluted to 2-8 nM in filtered SEC buffer (0.22 µm). Calibration was performed using a custom bAPO standard consisting of a mixture of apoferritin and β-amylase, spanning molecular weights of 56–506 kDa. Calibration quality was assessed by the R² value. For measurements, filtered buffer was first added to the wells for blank acquisition. Protein samples were then diluted directly on the coverslip to a final concentration of 2–10 nM. Data were acquired using Refeyn AcquireMP software and analyzed using Refeyn DiscoverMP.

### *In vitro* transcription of RNAs

Equimolar (33 μM) sense and antisense DNA strands (Ultramers, IDT), containing T7 polymerase promoter sequence (TAATACGACTCACTATA<u>GGG</u>) and -GGG included to increase transcriptional yield, were annealed in DNA annealing buffer (10 mM Tris–HCl pH 7.6, 50 mM NaCl and 1 mM EDTA pH 8), by incubating the DNA at 95°C for 3 min and slowly cool down to room temperature. RNA *in vitro* transcription was carried out with the HiScribe T7 High Yield RNA Synthesis Kit (NEB, E2040S) using 2 μg of annealed dsDNA template, 10 mM NTPs, and 1.5 μL of T7 polymerase mix in 1x reaction buffer, in a 20 μL reaction. The reaction was incubated at 37°C for 16 hours; DNA templates were degraded with 4 U of Turbo DNase (Invitrogen, AM2238) for 1 h at 37°C. RNA was isolated from each reaction with 400 μL of TRIzol LS reagent (Invitrogen) and 200 μL of 1-bromo-3-chloropropane (Sigma, B9673-200ML), and then again with 400 μL of 1-bromo-3-chloropropane by collecting the top aqueous phase. RNA was then precipitated in 1 volume of isopropanol and 0.75 M ammonium acetate overnight at −20°C and washed with 100% ethanol, followed by a 70% ethanol wash. RNA pellets were air-dried and resuspended in DEPC-treated water and stored at −80 °C. All *in vitro* transcribed RNA sequences are described in the Supplementary Table S3, and their quality can be assessed in Supplementary Data Fig.6.

### Electrophoretic mobility shift assay (EMSA)

Purified 6xHis-NSUN2 WT and NSUN2^C271S^-TwinStrep were buffer-exchanged into complex formation buffer (50 mM HEPES pH 7.0, 150 mM NaCl, 5 mM MgCl₂, 1 mM TCEP) using 0.5 mL Pierce protein concentrators (50,000 MWCO). *In vitro* transcribed RNA (1 μM) was unfolded by heating to 95°C for 2 minutes and then refolded for 20 minutes at 37 °C in RNA folding buffer (110 mM HEPES pH 8.0, 110 mM NaCl, 10 mM MgCl_2_). Folded RNA was incubated with 6xHis-NSUN2^WT^ or NSUN2^C271S^-TwinStrep (0, 0.2, 0.5, 1, 2, 5,10 μM) in binding buffer (50 mM HEPES pH 7.0, 150 mM NaCl, 5 mM MgCl2, 1 mM TCEP) for 30 minutes on ice. RNA-protein complexes were mixed with 1x Native gel loading dye (Invitrogen, AM8556) prior to separation on polyacrylamide TBE- PAGE gel (6%) in 1xTBE at 150V for 60 minutes and 4°C. RNA was visualized with a UV transillumination in a Gel Doc™ XR+ imager after staining with SYBR™ Gold. To estimate the binding affinity (*K_d_*), titration curves were obtained by quantification of free RNA bands in ImageJ. The percentage of bound RNA values was obtained by subtracting free RNA and then normalized to levels at 0 μM protein. The binding curves were fitted and binding parameters estimated using the specific binding Hill slope model in GraphPad Prism. Protein was visualized with Instant Blue Coomassie (Abcam) staining.

### Covalent adduct formation assay

*In vitro* transcribed RNA was unfolded by heating at 95 °C for 2 min and refolded by incubation at room temperature for 15 min in complex formation buffer (50 mM HEPES pH 7.0, 150 mM NaCl, 5 mM MgCl_2_, 1 mM TCEP). NSUN2^C271S^ was incubated with RNA and S-adenosyl-L-methionine (SAM; NEB) at protein to RNA and protein to SAM molar ratios of 1:1 or 1:2 and 1:50, respectively, for 30 min at 37 °C. Complexes were then mixed with Laemmli buffer, heated at 95 °C for 5 minutes, and run on SDS-PAGE (4-15%)(Biorad 456-1083) at 180V for 60 minutes. Bands were visualized by Instant Blue Coomassie (Abcam) staining.

### pXL-EM sample preparation

Purified NSUN2^C271S^ (80 µM) was buffer-exchanged into complex formation buffer (50 mM HEPES pH 7.0, 150 mM NaCl, 5 mM MgCl₂, 1 mM TCEP) using Amicon Ultra 4 mL concentrators (3,000 MWCO). Protein concentration was determined using a NanoDrop™ OneC spectrophotometer. Human tRNA^Asp^_GUC_ was unfolded by heating at 95 °C for 2–3 min and refolded by incubation at room temperature for 15 min in complex formation buffer. NSUN2^C271S^ was incubated with tRNA^Asp^_GUC_ and S-adenosyl-L-methionine (SAM; NEB) at protein to tRNA and protein to SAM molar ratios of 1:20 and 1:50, respectively, for 30 min at 37 °C.

For the crosslinked pXL-EM sample, sulfo-NHS-diazirine (sulfo-SDA) was dissolved in protein buffer to a final concentration of 6.5 µg/µL (1:0.38 protein to cross-linker weight/weight ratio). Crosslinking was initiated by adding 2 µL sulfo-SDA solution per sample and incubating for 30 min at room temperature in the dark. Samples were subsequently irradiated with UV light at 365 nm for 20 min to activate crosslinking. Reactions were quenched by the addition of 50 mM end concentration of Tris-HCl pH 7.0. In comparison, this step has been skipped for the non-crosslinked EM sample preparation.

Complex formation was assessed by SEC on a Superdex 200 Increase 10/300 column (Cytiva) equilibrated with SEC buffer and eluted with 1.5 CV. Purified non-crosslinked and sSDA-crosslinked samples were analyzed by SDS–PAGE using 3–8% Tris-acetate gels (Thermo Fisher Scientific) in 1× Tris-acetate running buffer. Thermo Scientific PageRuler Plus Prestained Protein Ladder (10–250 kDa) was loaded (3 µL) as a molecular weight marker. Electrophoresis was performed using the standard Tris-acetate program (150 V for 50 min). Bands were visualized by Instant Blue Coomassie staining (Abcam). Complex-containing fractions were concentrated to A280 ∼1 using Amicon Ultra concentrators (10,000 MWCO) by centrifugation at 3,500 × g at 4 °C to be ready for further cryoEM plunge-freezing.

Quantifoil R 1.2/1.3 Cu 300-mesh grids (Quantifoil Micro Tools) were glow-discharged for 60 s at 30 mA immediately before use. 3 μL of concentrated SEC sample was applied to grids on a Vitrobot Mark IV (Thermo Fisher Scientific), which was equilibrated to 4 °C and 100% relative humidity. For the non-crosslinked specimen, after 30 s of incubation, grids were blotted for 2 s (blot force set to 0, considering the calibration of our instrument), then plunge-frozen into liquid ethane. Grids were stored in liquid nitrogen until data acquisition. For the pXL-EM crosslinked specimen, after applying the sample, grids were blotted for 4 s (blot force set to 0, considering the calibration of our instrument), then plunge-frozen into liquid ethane. Grids were stored in liquid nitrogen until data acquisition.

### Cryo-EM data collection

Cryo-EM data were collected on a Titan Krios G4 microscope (Thermo Fisher Scientific) operating at 300 kV and equipped with a post-column Selectris X (Thermo Scientific) energy filter and a Falcon 4i (Thermo Scientific) direct electron detector. Movies were acquired in aberration-free image-shift (AFIS) mode at a nominal magnification of 215,000×, corresponding to a calibrated pixel size of 0.582 Å at the specimen level. Each movie was recorded with a total electron exposure of 60 e⁻/Å² equally distributed across multiple frames. For the non-crosslinked complex, a total of 11,268 movies were collected over a nominal defocus range of −0.5 to −1.75 µm (set points: −0.5, −0.75, −1.0, −1.25, −1.5, and −1.75 µm). For the pXL-EM crosslinked complex, a total of 31,517 movies were collected over a nominal defocus range of −0.8 to −1.8 µm (set points: −0.8, −1.0, −1.2, −1.4, −1.6, and −1.8 µm).

### Cryo-EM data processing

Both EM datasets have been processed in cryoSPARC v4.7.1 ^45^. The details of the EM processing are described in Supplementary Data Fig.2 for the pXL-EM complex and Supplementary Data Fig.3 for the non-crosslinked complex.

#### Non-crosslinked complex

The cryo-EM processing workflow is summarized in Supplementary Data Fig.3. For cryo-EM analysis of the non-crosslinked complex, 11,268 movie stacks were imported and processed in cryoSPARC v4.7.1 ^45^. Motion correction was performed using Patch Motion Correction with dose weighting, followed by Patch-based CTF estimation and Micrograph Curation, from which micrographs were selected based on CTF fit quality (threshold < 8 Å), resulting in a final dataset of 11,065 micrographs.

Initial particle picking was performed using the default blob picker with a particle diameter range of 120–180 Å, templates low-pass filtered to 10 Å, and a minimum separation distance of 0.75. Particles were extracted with a box size of 260 pixels and binned by a factor of 2. After initial particle clean up in 2D, ab initio reconstruction was performed, and particles were cleaned up with Heterogeneous Refinement. The cleanest from this first round of picking resulted in a Non-Uniform Refinement volume strongly affected by anisotropy. This particle set was used to train a TOPAZ neural network particle picker ^46^. Particle picking and refinement improvement proceeded through three iterative cycles, each consisting of: (i) TOPAZ particle picking and extraction, (ii) multiple rounds of 2D classification, (iii) Ab-Initio reconstruction, (iv) Heterogeneous Refinement, and (v) Non-Uniform Refinement of the best particle subsets.

The final round of TOPAZ training and picking yielded 2,133,046 particles, which were extracted with a box size of 260 pixels and binned by a factor of 2. These particles were subjected to a broad Heterogeneous Refinement using a plotting B-factor of 0 and a spherical mask diameter of 100 Å, with five classes generated from the best Non-Uniform Refinement volume together with previous volumes corresponding to noise or low-resolution classes observed during the iterative cleaning process.

Particles were subsequently re-extracted from the 11,063 micrographs using a box size of 260 pixels without Fourier-cropping. Non-Uniform Refinement in cryoSPARC was performed with per-particle scale minimisation enabled and an initial low-pass filter of 8 Å, using 708,339 particles, resulting in a reconstruction at 3.4 Å resolution with markedly reduced anisotropy compared to the final reconstruction of the previous rounds of picking. Preferential orientation of the reconstruction was assessed using the cFAR value.

To further improve particle homogeneity, 3D classification was performed on the 708,339 particles into five classes using a solvent mask, a filter resolution of 4 Å, and an initial low-pass filter of 8 Å. The best subset, comprising 183,116 unbinned particles, was refined using Non-Uniform Refinement with per-particle scale minimisation enabled, yielding a reconstruction at 3.1 Å resolution and a cFAR of 0.41.

Subsequently, Global CTF Refinement was performed to estimate anisotropic magnification, spherical aberration, and tetrafoil aberration, followed by Local CTF Refinement. Particles then underwent Reference-Based Motion Correction in cryoSPARC, followed by another round of Non-Uniform Refinement. The final Local Refinement yielded a reconstruction at 3.0 Å resolution, based on the 0.143 Fourier Shell Correlation (FSC) criterion (Extended Data Fig. 1). To generate the sharpened map, the cryoSPARC Sharpening Tool was used with standard settings and a B-factor of −40.

To resolve the continuous heterogeneity of the C-terminal region of NSUN2, 3D classification was performed using a focused mask on the PUA domain and tRNA acceptor stem. The classification was performed in input mode, tuning filter resolution to 4 Å, lowpass filtering the initial volumes to 8 Å. Three particle subsets obtained were independently refined via Non-Uniform Refinement using a solvent mask, followed by local refinement with the C-terminal Mask. The resulting volumes from the Non-Uniform Refinement and Local Refinement were combined to generate a composite map for each group, dissecting the continuous heterogeneity of the NSUN2 PUA domain and tRNA acceptor stem in three states.

#### pXL-EM complex

The cryo-EM processing workflow is summarized in Supplementary Data Fig. 2. For cryo-EM analysis of the pXL-EM complex, 31,511 movie stacks were imported and processed in cryoSPARC v4.7.1 ^45^. Motion correction was performed using Patch Motion Correction with dose weighting, followed by Patch-based CTF estimation. Micrographs were curated based on CTF fit quality (threshold < 10 Å) and Total full-frame motion distance (threshold < 60 pixels), resulting in 29,295 micrographs for further processing.

Initial particle picking was performed manually on a subset of 97 micrographs to generate a training set for the TOPAZ neural network particle picker ^46^. The model was trained iteratively over two rounds using particles selected after 2D classification and heterogeneous refinement. A starting pool of 762,007 particles was extracted with a box size of 420 pixels and Fourier-cropped to 210 pixels. The full set of particles was used to generate multiple initial volumes with Ab-Initio reconstruction, necessary to sort the heterogeneity of the sample. Iterative 2D classification yielded 182,794 particles, which were used to generate, after Heterogeneous Refinement and Non-Uniform Refinement, a volume of the complex with a resolution of 3.6 Å. This volume was used as input for 3D classification of 112,150 particles sorted via Heterogeneous Refinement from 461,308 particles cleaned via 2D classification. The 3D Classification into four classes was performed using a solvent mask, a filter resolution of 3 Å, and an initial low-pass filter of 8 Å. Following, the 33,780 particles classified in the best volume were extracted without Fourier-cropping. After Non-Uniform and Local Refinement, the particle set yielded a 3.5 Å resolution reconstruction.

Subsequently, Global CTF Refinement was performed to estimate anisotropic magnification, spherical aberration, and tetrafoil aberration, followed by Local CTF Refinement. Following, Non-Uniform and Local Refinement with the same parameters was performed.

These procedures resulted in a final reconstruction at 3.1 Å resolution, based on the 0.143 Fourier Shell Correlation (FSC) criterion (Fig.1, Extended Data Fig. 1). To generate the sharpened map, the cryoSPARC Sharpening Tool was used with standard settings and a B-factor of −15.

### Model building

Initial atomic models for NSUN2 and the tRNA^Asp^_GUC_ were generated independently using AlphaFold 3.0 (Supplementary Data Fig.1a,c) ^47^. IUPred2A was used to identify intrinsically disordered regions within the protein sequence (Supplementary Data Fig. 1b). The models were docked into the cryo-EM density using UCSF ChimeraX 1.10 and manually adjusted in Coot 0.9.8.96 to optimize the fit to the map. Iterative rounds of manual rebuilding and correction of geometry/rotamer outliers were performed in Coot prior to real-space refinement in Phenix 1.21.2 ^48,49^.

To model the catalytic covalent intermediate, an initial CIF restraint file for 5-methylcytosine (m^5^C) was generated using eLBOW (Phenix 1.21.2) ^50^ using the Chemical Component three-letter code 5MC, and subsequently manually modified to describe the covalent bond between m^5^C48 and the catalytic residue Cys321 of NSUN2. Bond lengths and angles were defined according to the established reaction chemistry of RNA cytosine-5 methyltransferases, in which the catalytic cysteine performs nucleophilic attack at the C6 position of cytosine to form a transient covalent intermediate during methyl transfer. The custom restraints were applied during refinement using the Phenix real-space refinement tool from the graphical interface, with the Coot-refined model used as the starting model. This enabled the formation of the covalent bond while maintaining appropriate stereochemistry throughout refinement, yielding the final refined structure. Validation was performed using Molprobity ^51^ and the wwPDB Validation Service (https://validate.wwpdb.org). The refinement table is attached in the Supplementary Table S1. Normal mode flexibility analysis was carried out with an elastic network model by elNemo webserver (http://www.elnemo.org/) ^52^.

### Protein-RNA UV-crosslinking Mass Spectrometry (MS) sample preparation

Purified NSUN2^C271S^ (∼3 mg/mL; ∼34 µM) was prepared in buffer containing 50 mM HEPES (pH 7.5), 150 mM NaCl, 5 mM MgCl₂, 1 mM TCEP, and 5% (v/v) glycerol. The NSUN2–tRNA complex was assembled at a molar ratio of NSUN2:tRNA:SAM = 1:2:10 by sequential addition of human tRNA^Asp^_GUC_ (final concentration 60 µM) and S-adenosyl-L-methionine (SAM; final concentration 300 µM), followed by addition of NSUN2 to yield a final protein concentration of ∼30 µM. Prior to complex assembly, the tRNA^Asp^_GUC_ was denatured for 3 min at 95 °C and immediately placed on ice. The complex was incubated for 60 min at 37 °C.

Following incubation, the sample was loaded onto a Superdex 200 Increase 10/300 column equilibrated in SEC buffer consisting of 50 mM HEPES (pH 7.0), 150 mM NaCl, 5 mM MgCl₂, and 0.5 mM TCEP. A total volume of 450 µL was injected and separated at a flow rate of 0.375 mL/min. Peak fractions corresponding to the first SEC peak were pooled (∼1 mL total), concentrated using a 3 kDa MWCO Amicon centrifugal concentrator, and used for downstream UV crosslinking. Complex quality was assessed by SDS–PAGE and mass photometry.

For UV-induced protein-RNA crosslinking, 50 µL aliquots of NSUN2-tRNA complex were placed in Protein LoBind tube lids and irradiated on ice at 254 nm using a total energy of 3.2 J/cm² (3200 mJ/cm²). After irradiation, samples were recovered into a single tube. As a negative control, an apo NSUN2^C271S^ sample has been treated identically.

Crosslinked samples were precipitated by adding 1/10 volume of 3 M sodium acetate (pH 5.1) and 3 volumes of ice-cold ethanol, followed by incubation at −20 °C for at least 2 h. Samples were centrifuged at 16,000 × g for 30 min at 4 °C. Pellets were washed with 80% ethanol and centrifuged again at 16,000 × g for 5–10 min at 4 °C. Pellets were resuspended in 25 µL 50 mM Tris-HCl (pH 8.0) containing 4 M urea, and diluted with 75 µL 50 mM Tris-HCl (pH 8.0) to reduce the urea concentration to 1 M.

RNA was digested using RNase A (Roche) at a nominal ratio of 5 µg RNase A per mg of protein sample. In this experiment, ∼900 μg of the 1:1 stoichiometry and 330 μg of the apo protein were processed. RNase A (ThermoFischer) was diluted 10-fold in 50 mM Tris-HCl (pH 8.0) and added to each sample following the previous nominal ratio. Digestion was performed for 2 h at 52 °C with shaking at 300 rpm. Trypsin digestion was performed overnight at 37 °C using a 24:1 protein-to-enzyme ratio, corresponding to ∼2 µg trypsin per 50 µg sample. Trypsin activity was subsequently quenched by incubation at 70 °C for 10 min. Peptides were desalted using C18 StageTips (CDS Analytical) and dried by vacuum centrifugation (SpeedVac; 45 °C; ∼40 min). RNA–peptide crosslinks were TiO2-enriched using the High Select™ Phosphopeptide Enrichment kit (Thermo Fischer Scientific, A32993) following the manufacturer’s instructions.

### Protein-RNA UV-crosslinking MS acquisition

Protein-RNA cross-linked samples after TiO_2_ enrichment were injected for liquid chromatography-mass spectrometry (LC-MS) acquisition. The LC-MS platform consisted of a Vanquish Neo system (Thermo Fisher Scientific) connected to an Orbitrap Eclipse Tribrid mass spectrometer (Thermo Fisher Scientific) operating under Tune 4.4321. Mobile phases consisted of 0.1% v/v formic acid in water (mobile phase A) and 0.1% formic acid in 80% acetonitrile/water v/v (mobile phase B). Samples were dissolved in mobile phase A. Samples were separated on an EASY-Spray PepMap Neo column (75 µm x 50 cm) (ThermoFisher Scientific). Peptides were separated at a flow rate of 300 nL/min using a linear 96 min gradient of 5-95% B, followed by a 10 min column wash at 95%B, and column equilibration. MS1 spectra (scan range 400-1600 m/z) were acquired with a resolution of 120,000 and an automated gain control target set to standard, with a maximum injection time of 50 ms. Cycle time was set to 3 seconds. The source RF lens was set to 35%. Dynamic exclusion was set to 30 seconds. A precursor charge filter with z=2-7 was adopted. Precursors were selected based on a data-dependent decision strategy and subjected to stepped higher-energy collisional dissociation (HCD) fragmentation with normalized collision energies of 18, 23, and 28. MS2 scans were acquired with a normalized gain control target of 250%. The maximum injection time was set to 250 ms, and the Orbitrap resolution to 60,000. Fluoranthene was used for lock mass correction during MS runs.

### Protein-RNA UV-crosslinking MS data analysis

Raw files were analyzed using the OpenMS-NuXL ^53^ stand-alone tool via TOPPAS ^54^. The OpenNuXL node was configured with the following settings: precursor mass tolerance: 3 ppm, fragment mass tolerance: 10 ppm, min_charge 2, max_charge 5, isotope correction disabled, modifications: variable – methionine oxidation, with a maximum of 2 variable modifications per peptide, enzyme: Trypsin/P, missed_cleavages: 2. A fasta file containing the sequence of NSUN2^C271S^-TwinStrep and common mass spec contaminants was used for analysis. The RNA-UV (UGCA) RNPxl preset was used for the analysis, with a maximum oligonucleotide length set to 5. All the other parameters were set to the default. Additionally, a FragPipe 23.-0 ^55^ open modification search ^56^ was performed using suggested parameters ^57^ to corroborate all the identified peptide-RNA adducts. A final table containing all the identified OpenMS-NuXL peptide-RNA pairs was exported and attached in Supplementary Data 1. Exemplary fragment ion spectra for some of the identified protein-RNA UV crosslinks are shown in Supplementary Data Fig.4.

### Chemical crosslinking MS sample preparation

Purified NSUN2^C271S^ was used for complex formation. The protein was stored in 50 mM HEPES (pH 7.5), 150 mM NaCl, 5 mM MgCl₂, 1 mM TCEP, and 5% (v/v) glycerol. Protein concentration was estimated from absorbance measurements, corresponding to ∼3 mg/mL (∼34 µM).

Complex formation reactions were assembled at a molar ratio of NSUN2:tRNA:SAM = 1:2:10, corresponding to final concentrations of 30 µM NSUN2, 60 µM human tRNA^Asp^_GUC_, and 300 µM S-adenosyl-L-methionine (SAM). The tRNA^Asp^_GUC_ was mixed with buffer and heat-treated for 3 min at 95 °C to unfold the RNA and promote refolding. After cooling on ice, NSUN2^C271S^ was added to the reaction mixture, followed by the addition of SAM. The reaction mixture was incubated for 60 min at 37 °C.

Sulfo-NHS-diazirine (sulfo-SDA) was dissolved in protein buffer to a final concentration of 6.5 µg/µL (1:0.38 protein to cross-linker weight/weight ratio). Crosslinking was initiated by adding 2 µL sulfo-SDA solution per sample and incubating for 30 min at room temperature, protected from light. Samples were subsequently irradiated with UV light at 365 nm for 20 min to activate crosslinking. Reactions were quenched by the addition of 50 mM end concentration of ammonium bicarbonate (ABC).

Crosslinked samples were analysed by SDS–PAGE using 3–8% Tris-acetate gels in 1× Tris-acetate running buffer. A total of ∼33 ug of protein was loaded per lane, for a total of 3 lanes, for a total of ∼100 ug of protein. Thermo Scientific PageRuler Plus Prestained Protein Ladder (10–250 kDa) was loaded (3 µL) as a molecular weight marker. Electrophoresis was performed using the standard Tris-acetate program (150 V for 50 min).

Gels were stained with Instant Blue Coomassie stain (Abcam) for 15 min and imaged using a gel documentation system. Bands corresponding to NSUN2–tRNA complexes were excised for downstream analysis. Gel pieces were dehydrated with 100% acetonitrile and stored at −20 °C until trypsin digestion and peptide cleanup. In-gel digestion of sulfo-SDA crosslinked samples was performed according to a previously published protocol ^58^. Shortly thereafter, gel pieces were destained using a solution containing 100% acetonitrile and 50 mM ABC, added in equal volumes, for 10-30 minutes, and mixed with a ThermoMixer at 900 r.p.m. Gel pieces were dehydrated with 100% acetonitrile between every washing step. Samples were reduced by incubation with 10 mM dithiothreitol (DTT) for 30 minutes at 37 °C, followed by alkylation with 55 mM iodoacetamide (IAA) for 30 minutes in the dark. After reduction/alkylation, gel pieces were dehydrated with acetonitrile, and 2 µg trypsin was added to each sample. Peptides were recovered using a shrink/expansion protocol using sequential addition of i) acetonitrile, ii) 0.1% trifluoroacetic acid, iii) acetonitrile. Samples were concentrated using a SpeedVac and finally desalted using C18 Stage Tips (CDS Analytical).

### Chemical crosslinking MS acquisition

Sulfo-SDA-crosslinked samples were injected for each liquid chromatography-mass spectrometry (LC-MS) acquisition. The LC-MS platform consisted of a Vanquish Neo system (Thermo Fisher Scientific) connected to an Orbitrap Eclipse Tribrid mass spectrometer (Thermo Fisher Scientific) equipped with a FAIMSpro device, operating under Tune 4.2.4321. Mobile phases consisted of 0.1% v/v formic acid in water (mobile phase A) and 0.1% formic acid in 80% acetonitrile/water v/v (mobile phase B). Samples were dissolved in mobile phase A. The FAIMSpro device was set to standard resolution with a carrier gas flow of 4.6L/min. The samples were separated on an EASY-Spray PepMap Neo column (75 µm × 50 cm) (Thermo Fisher Scientific).

MS1 spectra were acquired with a resolution of 120,000 and an automated gain control target set to standard, with a maximum injection time of 50 ms. Cycle time was set to 2.5 seconds. The source RF lens was set to 35%. Dynamic exclusion was set to 30 seconds per FAIMS control voltage or 60 seconds per experiment with a single FAIMS control voltage. A precursor charge filter was set to z=3-7. Precursors were selected based on a data-dependent decision tree strategy ^59^, prioritizing charge states 4-7 and subjected to stepped HCD fragmentation with normalized collision energies of 24, 27, and 30. MS2 scans were acquired with a normalized gain control target of 750%. The maximum injection time was set to 250 ms, and the Orbitrap resolution to 60,000. Four different FAIMS control voltages acquisitions were performed for the monomeric NSUN2 sulfoSDA crosslinked band, specifically at −45, −55, −65, and −40/-65.

### Chemical crosslinking MS data analysis

Raw files were converted to the mgf format using ProteoWizard MSconvert (version 3.0.24184-c8389fe) ^60^. A recalibration of the MS1 and MS2 data was conducted based on high-confidence (<1% false discovery rate) linear peptide identifications using the xiSEARCH preprocessing pipeline (accessible at https://github.com/Rappsilber-Laboratory/preprocessing). Crosslinking MS database search was performed in xiSEARCH (version 1.8.9) ^61^ on a database comprising the NSUN2 sequence and common mass spectrometry contaminants derived from MaxQuant ^62^, using trypsin as protease with 4 missed cleavages. Precursor mass error tolerance was set to 3 ppm and MS2 error tolerance to 5 ppm. Carbamidomethylation of cysteine was defined as a fixed modification. The search included asparagine deamidation as linear modification, methionine oxidation, SDA-loop link (+82.04186484 Da), and SDA-OH (+100.0524 Da) as variable modifications. The SDA crosslinker was defined as cleavable ^63^ and reactive from K or protein N-termini to any amino acid. The search was set to account for noncovalent gas-phase associations. Results were filtered to 1% FDR at the residue pair level using xiFDR 2.10 with the following prefilters: (i) fragment unique crosslinked matched conservative >1; (ii) peptide1 unique matched non lossy >1; (iii) peptide2 unique matched non lossy >1. A final table containing all the identified crosslinks was exported and attached in Supplementary Data 2.

### Molecular Dynamics Simulation

In the cryo-EM structure, the missing loops for protein (residue id 593-598 and 664-671) were copied from the structure: AF-Q08J23-F1-v6 from AlphaFold protein structure database ^64^. These added residues were energy minimized using the 3D builder in Maestro (Schrödinger *Schrödinger Release 2021-3: Maestro (2021)*. The input models for MD simulations were prepared using the CHARMM-GUI server ^65–68^. The N- and C-termini of the protein were capped with acetyl and N-methyl groups, respectively. The protonation states of amino acids at pH 7.4 were predicted using PROPKA calculations ^69,70^. The system was solvated in a cubic box of TIP3P ^71^ water molecules with dimensions 117 × 117 × 117 Å³, and periodic boundary conditions were applied. Na⁺ and Cl⁻ ions were added to achieve a final concentration of 0.15 M. After solvation and addition of salt ions, the system comprised approximately 150,000 atoms. Topology and parameter for S-adenosylhomocysteine (SAH) were created using the CHARMM-GUI server employing the CHARMM General Force Field ^72^. Minimization, equilibration, and production were carried out using the PMEMD program from the AMBER24 package ^73,74^ with the CHARMM36m force field ^75^. Initial energy minimization was carried out in two stages: first, 4000 steps of steepest descent minimization were applied, followed by 6000 steps of conjugate gradient minimization. The system was progressively heated to 310 K over 100 ps under NVT conditions, using a Langevin thermostat ^76^ with a friction coefficient of 2.0 ps⁻¹ during both equilibration and production phases. Equilibration was carried out in the NPT ensemble with pressure regulation and a 1 fs integration timestep. The procedure involved eight consecutive stages during which positional restraints on the nucleotides, protein, and the small molecule (SAH) were progressively reduced from 5 to 0.10 kcal·mol^-1^·Å^-2^. The final stage of equilibration involved a short 1 ns simulation carried out in the NPT ensemble with a 2 fs timestep. This initial segment was excluded from further analysis. Both equilibration and production runs were conducted under isotropic pressure coupling using the Berendsen barostat ^77^. A 9 Å cutoff was applied for nonbonded interactions, including van der Waals and electrostatics. Trajectory snapshots were recorded at 100 ps intervals, yielding 5000 frames over the 500 ns simulation period. Two independent simulations were performed for each system. Although each production run was initiated from the same equilibrated structure, distinct random seeds were assigned to the Langevin thermostat to ensure statistical independence without modifying the initial coordinates or velocities. Data from all replicates were incorporated into the subsequent analyses. Trajectory analyses included root-mean-square deviation (RMSD), root-mean-square fluctuation (RMSF), were performed using CPPTRAJ ^78^ from the AmberTools24 ^79^ suite.

## DATA AVAILABILITY

Cryo-EM density map (post-processed map, half maps and mask) have been deposited in the Electron Microscopy Data Bank under accession codes EMD-57136 (NSUN2^C271S^-tRNA^Asp^_GUC_ pXL-EM sample) and EMD-57155 (non-crosslinked sample). The coordinates have been deposited in the Protein Data Bank under accession numbers 29FB for the NSUN2^C271S^-tRNA^Asp^_GUC_ pXL-EM complex.

The sulfo-SDA crosslinking mass spectrometry data have been deposited to the ProteomeXchange^80^ Consortium via the PRIDE^81^ partner repository with the dataset identifier PXD075494. The UV-RNA crosslinking mass spectrometry data have been deposited with PRIDE identifier PXD075678. Code for mapping RNA-protein crosslinks identified by NuXL to the NSUN2^C271S^-tRNA^Asp^_GUC_ model is available at https://github.com/grandrea/RNA-protein-xl-map.

Unique materials generated in this study will be made available from the corresponding author upon reasonable request, subject to completion of a standard material transfer agreement.

## ACKNOWLEDGMENTS

We thank all members of the Casañal lab for helpful discussions. We thank S. Buscarini for help with optimisation of cloning, expression and purification of NSUN2(C271S); T. G. Martin for cryo-EM processing advice; the National Facility for Structural Biology at Human Technopole, in particular P. Swuec, G. D’Urso, and M. Cavinato, for cryo-EM technical support and assistance; P. Swuec for his contributions to the design of the pXL-EM workflow strategy; P.A. Scardua, and J. Weber for help with insect cell culture; Daniele Colombo (IT Department, Human Technopole) for software installation and computational assistance; and A. C. Innis, for advice on model building of covalent adducts.

This work was supported by Human Technopole (A.C.), an EMBO Long-Term Fellowship (ALTF 894-2022; E.C.L.), and by the Medical Research Council [BVR025900], EPA Trust Fund [BVR01670], and Lee Placito Fund (M.G.).

## AUTHOR CONTRIBUTIONS

A.C. conceived and supervised the project; E.C.L. contributed to project conception, led the experimental strategy, designed and performed most experiments. E.C.L., A.D.F., A.R., and J.R.V. cloned, expressed and purified protein complexes. E.C.L., M.L. A.R., and J.R.V. performed MP experiments. A.D.F. performed *in vitro* transcription, and EMSA experiments. E.C.L. and J.R.V. developed and optimized the covalent adduct formation assay. E.C.L. and M.L. prepared cryo-EM samples and grids, collected cryo-EM data, processed the data, and performed model building, refinement and structural interpretation. K.L. performed molecular dynamics simulations and analysed the data. E.C.L. and A.D.I. prepared samples for UV and chemical crosslinking MS experiments with assistance from A.G.; A.D.I. collected and analysed XL-MS data; A.G. wrote scripts for analysis of UV-crosslinking data. E.C.L. and M.L. prepared all illustrations with A.C. assistance, and contributions from A.D.F. M.G. contributed to the design and interpretation of the biochemical experiments, supervised A.D.F., and, together with A.C., helped shape the overall interpretation of the study. A.C., M.G., and E.C.L. wrote the manuscript; M.L. provided feedback, and all authors reviewed and approved the final version.

## CORRESPONDING AUTHORS

Correspondence to Monika Gullerova and/or Ana Casañal

## REFERENCES

1. Bohnsack, K. E., Höbartner, C. & Bohnsack, M. T. Eukaryotic 5-methylcytosine (m^5^C) RNA Methyltransferases: Mechanisms, Cellular Functions, and Links to Disease. Genes 10, (2019).

2. Jonkhout, N. et al. The RNA modification landscape in human disease. RNA 23, 1754–1769 (2017).

3. Frye, M., Harada, B. T., Behm, M. & He, C. RNA modifications modulate gene expression during development. Science 361, 1346–1349 (2018).

4. Cui, L. et al. RNA modifications: importance in immune cell biology and related diseases. Signal Transduct. Target. Ther. 7, 334 (2022).

5. Delaunay, S., Helm, M. & Frye, M. RNA modifications in physiology and disease: towards clinical applications. Nat. Rev. Genet. 25, 104–122 (2024).

6. Trixl, L. & Lusser, A. The dynamic RNA modification 5-methylcytosine and its emerging role as an epitranscriptomic mark. Wiley Interdiscip. Rev. RNA 10, e1510 (2019).

7. Lu, Y. et al. RNA 5-methylcytosine modification: Regulatory molecules, biological functions, and human diseases. Genomics Proteomics Bioinformatics 22, qzae063 (2024).

8. Moon, J., Lee, H., Jang, Y. & Kim, S.-K. NSUN-mediated m5C RNA modification in stem cell regulation. Cells 14, 1609 (2025).

9. Yakubovskaya, E. et al. Structure of the essential MTERF4:NSUN4 protein complex reveals how an MTERF protein collaborates to facilitate rRNA modification. Structure 20, 1940–1947 (2012).

10. Cipullo, M., Gesé, G. V., Khawaja, A., Hällberg, B. M. & Rorbach, J. Structural basis for late maturation steps of the human mitoribosomal large subunit. Nat. Commun. 12, 3673 (2021).

11. Lenarčič, T. et al. Stepwise maturation of the peptidyl transferase region of human mitoribosomes. Nat. Commun. 12, 3671 (2021).

12. Hillen, H. S. et al. Structural basis of GTPase-mediated mitochondrial ribosome biogenesis and recycling. Nat. Commun. 12, 3672 (2021).

13. Cheng, J., Berninghausen, O. & Beckmann, R. A distinct assembly pathway of the human 39S late pre-mitoribosome. Nat. Commun. 12, 4544 (2021).

14. Chandrasekaran, V. et al. Visualizing formation of the active site in the mitochondrial ribosome. Elife 10, (2021).

15. Nguyen, T. G., Ritter, C. & Kummer, E. Structural insights into the role of GTPBP10 in the RNA maturation of the mitoribosome. Nat. Commun. 14, 7991 (2023).

16. Zhong, F. et al. NSUN6 inhibitor discovery guided by its mRNA substrate bound crystal structure. Structure 33, 443–450.e4 (2025).

17. Liu, R.-J., Long, T., Li, J., Li, H. & Wang, E.-D. Structural basis for substrate binding and catalytic mechanism of a human RNA:m5C methyltransferase NSun6. Nucleic Acids Res. 45, 6684–6697 (2017).

18. Selmi, T. et al. Sequence- and structure-specific cytosine-5 mRNA methylation by NSUN6. Nucleic Acids Res. 49, 1006–1022 (2021).

19. Frye, M. & Watt, F. M. The RNA methyltransferase Misu (NSun2) mediates Myc-induced proliferation and is upregulated in tumors. Curr. Biol. 16, 971–981 (2006).

20. Blanco, S. et al. Aberrant methylation of tRNAs links cellular stress to neuro-developmental disorders. EMBO J. 33, 2020–2039 (2014).

21. Khan, M. A. et al. Mutation in NSUN2, which encodes an RNA methyltransferase, causes autosomal-recessive intellectual disability. Am. J. Hum. Genet. 90, 856–863 (2012).

22. Martinez, F. J. et al. Whole exome sequencing identifies a splicing mutation in NSUN2 as a cause of a Dubowitz-like syndrome. J. Med. Genet. 49, 380–385 (2012).

23. Sajini, A. A. et al. Loss of 5-methylcytosine alters the biogenesis of vault-derived small RNAs to coordinate epidermal differentiation. Nat. Commun. 10, 2550 (2019).

24. Sun, Z. et al. Aberrant NSUN2-mediated m5C modification of H19 lncRNA is associated with poor differentiation of hepatocellular carcinoma. Oncogene 39, 6906–6919 (2020).

25. Jiang, Y. et al. NSUN2-mediated RNA m5C modification drives multiple myeloma progression by enhancing the stability of HIP1 mRNA. Sci. Rep. 15, 27888 (2025).

26. Li, C., Yuan, Y., Jiang, X. & Wang, Q. Roles and mechanisms of NSUN2-mediated RNA m5C modification in cancer progression and immune modulation. Front. Immunol. 16, 1702436 (2025).

27. Tuorto, F. et al. RNA cytosine methylation by Dnmt2 and NSun2 promotes tRNA stability and protein synthesis. Nat. Struct. Mol. Biol. 19, 900–905 (2012).

28. Shinoda, S. et al. Mammalian NSUN2 introduces 5-methylcytidines into mitochondrial tRNAs. Nucleic Acids Res. 47, 8734–8745 (2019).

29. Liu, Y. & Santi, D. V. m5C RNA and m5C DNA methyl transferases use different cysteine residues as catalysts. Proc. Natl. Acad. Sci. U. S. A. 97, 8263–8265 (2000).

30. King, M. Y. & Redman, K. L. RNA methyltransferases utilize two cysteine residues in the formation of 5-methylcytosine. Biochemistry 41, 11218–11225 (2002).

31. Redman, K. L. Assembly of protein-RNA complexes using natural RNA and mutant forms of an RNA cytosine methyltransferase. Biomacromolecules 7, 3321–3326 (2006).

32. Squires, J. E. et al. Widespread occurrence of 5-methylcytosine in human coding and non-coding RNA. Nucleic Acids Res. 40, 5023–5033 (2012).

33. Alessandro, B., Chamera, S., Cecatiello, V., Andrecka, J. & Vannini, A. Cryo-EM and single molecule visualization unravel the role of human condensin II activation by M18BP1 in driving DNA compaction. bioRxiv 2026.02.02.703226 (2026) doi:10.64898/2026.02.02.703226.

34. Biela, A. et al. The diverse structural modes of tRNA binding and recognition. J. Biol. Chem. 299, 104966 (2023).

35. Abbasi-Moheb, L. et al. Mutations in NSUN2 cause autosomal-recessive intellectual disability. Am. J. Hum. Genet. 90, 847–855 (2012).

36. Gonskikh, Y. et al. Spatial regulation of NSUN2-mediated tRNA m5C installation in cognitive function. Nucleic Acids Res. 53, (2025).

37. Li, J. et al. Structural basis of regulated m7G tRNA modification by METTL1-WDR4. Nature 613, 391–397 (2023).

38. Corbeski, I. et al. The catalytic mechanism of the RNA methyltransferase METTL3. Elife 12, (2024).

39. Ruiz-Arroyo, V. M. et al. Structures and mechanisms of tRNA methylation by METTL1-WDR4. Nature 613, 383–390 (2023).

40. Tao, Y. et al. Chemical proteomic discovery of isotype-selective covalent inhibitors of the RNA methyltransferase NSUN2. Angew. Chem. Int. Ed Engl. 62, e202311924 (2023).

41. Yu, S., Peng, Q., Wei, W., Li, X. & Long, S. AI-driven virtual screening platform identifies novel NSUN2 inhibitor candidates for targeted cancer therapy: a computational drug discovery approach. NPJ Precis. Oncol. 1–22 (2026).

42. Chen, T. et al. NSUN2 is a glucose sensor suppressing cGAS/STING to maintain tumorigenesis and immunotherapy resistance. Cell Metab. 35, 1782–1798.e8 (2023).

43. Miao, W. et al. Glucose binds and activates NSUN2 to promote translation and epidermal differentiation. Nucleic Acids Res. 52, 13577–13593 (2024).

## REFERENCES (Materials & Methods)

44. Bieniossek, C., Richmond, T. J. & Berger, I. MultiBac: multigene baculovirus-based eukaryotic protein complex production. Curr. Protoc. Protein Sci. **Chapter** 5, Unit 5.20 (2008).

45. Punjani, A., Rubinstein, J. L., Fleet, D. J. & Brubaker, M. A. cryoSPARC: algorithms for rapid unsupervised cryo-EM structure determination. Nat. Methods 14, 290–296 (2017).

46. Bepler, T. et al. Positive-unlabeled convolutional neural networks for particle picking in cryo-electron micrographs. Nat. Methods 16, 1153–1160 (2019).

47. Abramson, J. et al. Accurate structure prediction of biomolecular interactions with AlphaFold 3. Nature 630, 493–500 (2024).

48. Liebschner, D. et al. Macromolecular structure determination using X-rays, neutrons and electrons: recent developments in Phenix. Acta Crystallogr. D Struct. Biol. 75, 861–877 (2019).

49. Afonine, P. V. et al. New tools for the analysis and validation of cryo-EM maps and atomic models. Acta Crystallogr. D Struct. Biol. 74, 814–840 (2018).

50. Moriarty, N. W., Grosse-Kunstleve, R. W. & Adams, P. D. electronic Ligand Builder and Optimization Workbench (eLBOW): a tool for ligand coordinate and restraint generation. Acta Crystallogr. D Biol. Crystallogr. 65, 1074–1080 (2009).

51. Williams, C. J. et al. MolProbity: More and better reference data for improved all-atom structure validation. Protein Sci. 27, 293–315 (2018).

52. Suhre, K. & Sanejouand, Y.-H. ElNemo: a normal mode web server for protein movement analysis and the generation of templates for molecular replacement. Nucleic Acids Res. 32, W610–4 (2004).

53. Welp, L. M. et al. Chemical crosslinking extends and complements UV crosslinking in analysis of RNA/DNA nucleic acid-protein interaction sites by mass spectrometry. Nucleic Acids Res. 53, gkaf727 (2025).

54. Junker, J., et al. TOPPAS: a graphical workflow editor for the analysis of high-throughput proteomics data. J. Proteome Res. 11, 3914–3920 (2012).

55. Kong, A. T., Leprevost, F. V., Avtonomov, D. M., Mellacheruvu, D. & Nesvizhskii, A. I. MSFragger: ultrafast and comprehensive peptide identification in mass spectrometry-based proteomics. Nat. Methods 14, 513–520 (2017).

56. Yu, F. et al. Identification of modified peptides using localization-aware open search. Nat. Commun. 11, 4065 (2020).

57. Sarnowski, C. P. et al. A highly sensitive protein-RNA cross-linking mass spectrometry workflow with enhanced structural modeling potential. Nucleic Acids Res. 53, gkaf523 (2025).

58. Shevchenko, A., Tomas, H., Havlis, J., Olsen, J. V. & Mann, M. In-gel digestion for mass spectrometric characterization of proteins and proteomes. Nat. Protoc. 1, 2856–2860 (2006).

59. Kolbowski, L., Mendes, M. L. & Rappsilber, J. Optimizing the parameters governing the fragmentation of cross-linked peptides in a tribrid mass spectrometer. Anal. Chem. 89, 5311–5318 (2017).

60. Chambers, M. C. et al. A cross-platform toolkit for mass spectrometry and proteomics. Nat. Biotechnol. 30, 918–920 (2012).

61. Mendes, M. L. et al. An integrated workflow for crosslinking mass spectrometry. Mol. Syst. Biol. 15, e8994 (2019).

62. Cox, J. & Mann, M. MaxQuant enables high peptide identification rates, individualized p.p.b.-range mass accuracies and proteome-wide protein quantification. Nat. Biotechnol. 26, 1367–1372 (2008).

63. Iacobucci, C. et al. Carboxyl-photo-reactive MS-cleavable cross-linkers: Unveiling a hidden aspect of diazirine-based reagents. Anal. Chem. 90, 2805–2809 (2018).

64. Fleming, J. et al. AlphaFold Protein Structure Database and 3D-Beacons: New data and capabilities. J. Mol. Biol. 437, 168967 (2025).

65. Brooks, B. R. et al. CHARMM: the biomolecular simulation program. J. Comput. Chem. 30, 1545–1614 (2009).

66. Jo, S., Kim, T., Iyer, V. G. & Im, W. CHARMM-GUI: a web-based graphical user interface for CHARMM. J. Comput. Chem. 29, 1859–1865 (2008).

67. Lee, J. et al. CHARMM-GUI Input Generator for NAMD, Gromacs, Amber, Openmm, and CHARMM/OpenMM Simulations using the CHARMM36 Additive Force Field. Biophys. J. 110, 641a (2016).

68. Lee, J. et al. CHARMM-GUI supports the Amber force fields. J. Chem. Phys. 153, 035103 (2020).

69. Olsson, M. H. M., Søndergaard, C. R., Rostkowski, M. & Jensen, J. H. PROPKA3: Consistent treatment of internal and surface residues in empirical pKa predictions. J. Chem. Theory Comput. 7, 525–537 (2011).

70. Søndergaard, C. R., Olsson, M. H. M., Rostkowski, M. & Jensen, J. H. Improved treatment of ligands and coupling effects in empirical calculation and rationalization of pKa values. J. Chem. Theory Comput. 7, 2284–2295 (2011).

71. Mark, P. & Nilsson, L. Structure and dynamics of the TIP3P, SPC, and SPC/E water models at 298 K. J. Phys. Chem. A 105, 9954–9960 (2001).

72. Vanommeslaeghe, K. et al. CHARMM general force field: A force field for drug-like molecules compatible with the CHARMM all-atom additive biological force fields. J. Comput. Chem. 31, 671–690 (2010).

73. Case, D. A. et al. Recent developments in Amber biomolecular simulations. J. Chem. Inf. Model. 65, 7835–7843 (2025).

74. Weiner, P. K. & Kollman, P. A. AMBER: Assisted model building with energy refinement. A general program for modeling molecules and their interactions. J. Comput. Chem. 2, 287–303 (1981).

75. Huang, J. et al. CHARMM36m: an improved force field for folded and intrinsically disordered proteins. Nat. Methods 14, 71–73 (2017).

76. Davidchack, R. L., Ouldridge, T. E. & Tretyakov, M. V. New Langevin and gradient thermostats for rigid body dynamics. J. Chem. Phys. 142, 144114 (2015).

77. Berendsen, H. J. C., Postma, J. P. M., van Gunsteren, W. F., DiNola, A. & Haak, J. R. Molecular dynamics with coupling to an external bath. J. Chem. Phys. 81, 3684–3690 (1984).

78. Roe, D. R., Cheatham, T. E. & Ptraj, I. III PTRAJ and CPPTRAJ: Software for Processing and Analysis of Molecular Dynamics Trajectory Data. Journal of Chemical Theory and Computation 9, 3084–3095 (2013).

79. Case, D. A. et al. AmberTools. J. Chem. Inf. Model. 63, 6183–6191 (2023).

80. Deutsch, E. W. et al. The ProteomeXchange consortium in 2026: making proteomics data FAIR. Nucleic Acids Res. 54, D459–D469 (2026).

81. Perez-Riverol, Y. et al. The PRIDE database at 20 years: 2025 update. Nucleic Acids Res. 53, D543–D553 (2025).

