## Supplementary figures for "Molecular basis of tRNA modification by the human m^5^C methyltransferase NSUN2"

### EXTENDED DATA (with legends)

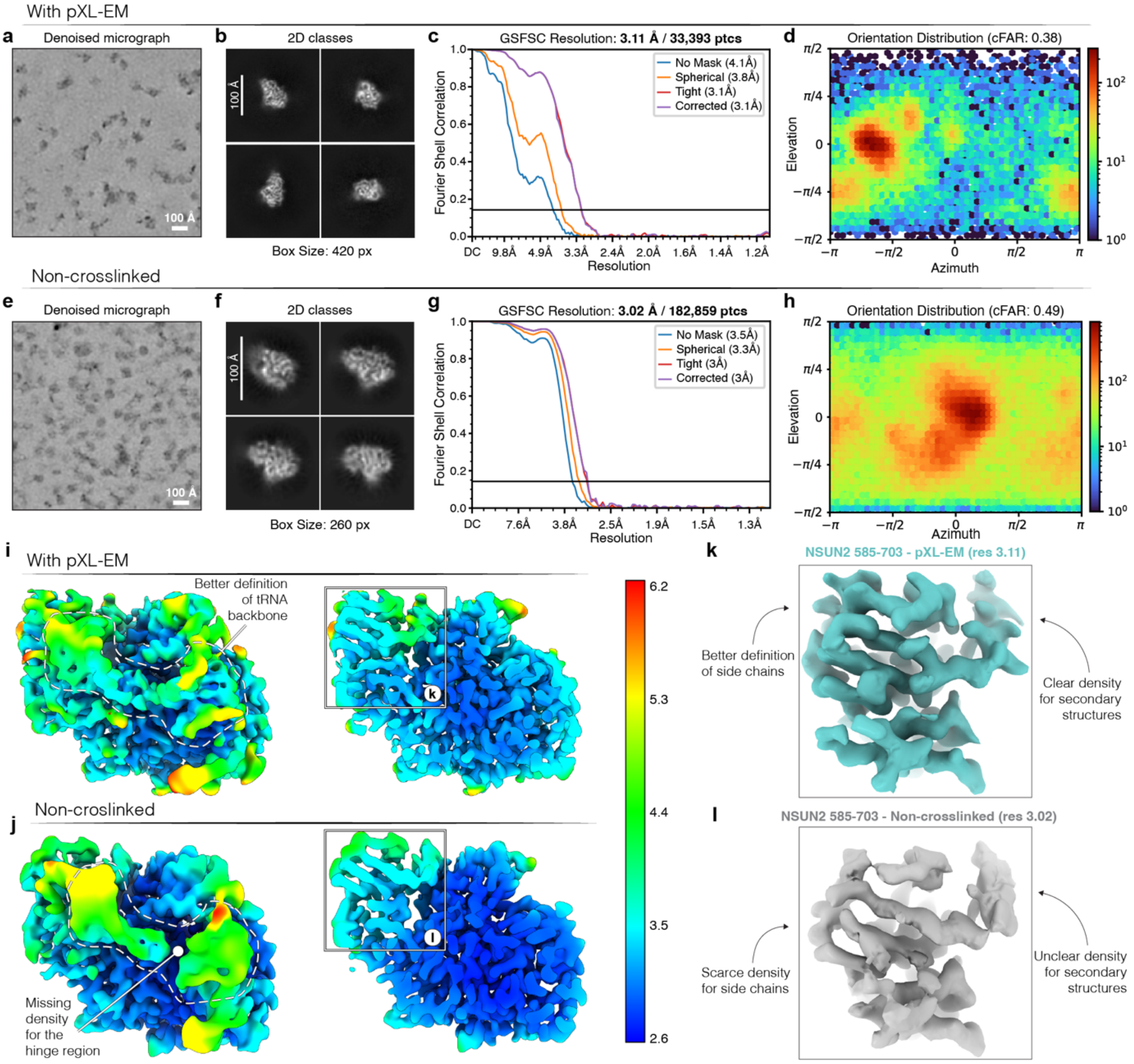

**Extended Data Fig 1 | Cryo-EM data quality for NSUN2-tRNA$^{Asp}_{GUC}$ with and without pXL-EM stabilisation.**

**a-d**, Data collected with pXL-EM (sulfo-SDA photo-crosslinking). **a**, Representative denoised micrograph. **b**, Representative 2D class averages. **c**, Gold-standard Fourier shell correlation (GSFSC) curve between independently refined half-maps, with the nominal resolution estimated at FSC = 0.143. **d**, Viewing direction distribution (cryoSPARC) with cFAR indicated.

**e-h**, Data collected in the absence of crosslinker conditions. **e**, Representative denoised micrograph. **f**, Representative 2D class averages. **g**, Gold-standard FSC curve and nominal resolution estimate (FSC = 0.143). **h**, Viewing direction distribution (cryoSPARC) with cFAR indicated.

**i,j**, Local-resolution maps for the final reconstructions (colour scale, Å), highlighting improved definition of the tRNA backbone in the pXL-EM dataset and reduced density in flexible regions (including the hinge region) in the non-crosslinked dataset.

**k,l**, Representative map features for NSUN2 residues 585-703, showing improved density for secondary structure and side chains in the pXL-EM reconstruction relative to the non-crosslinked reconstruction.

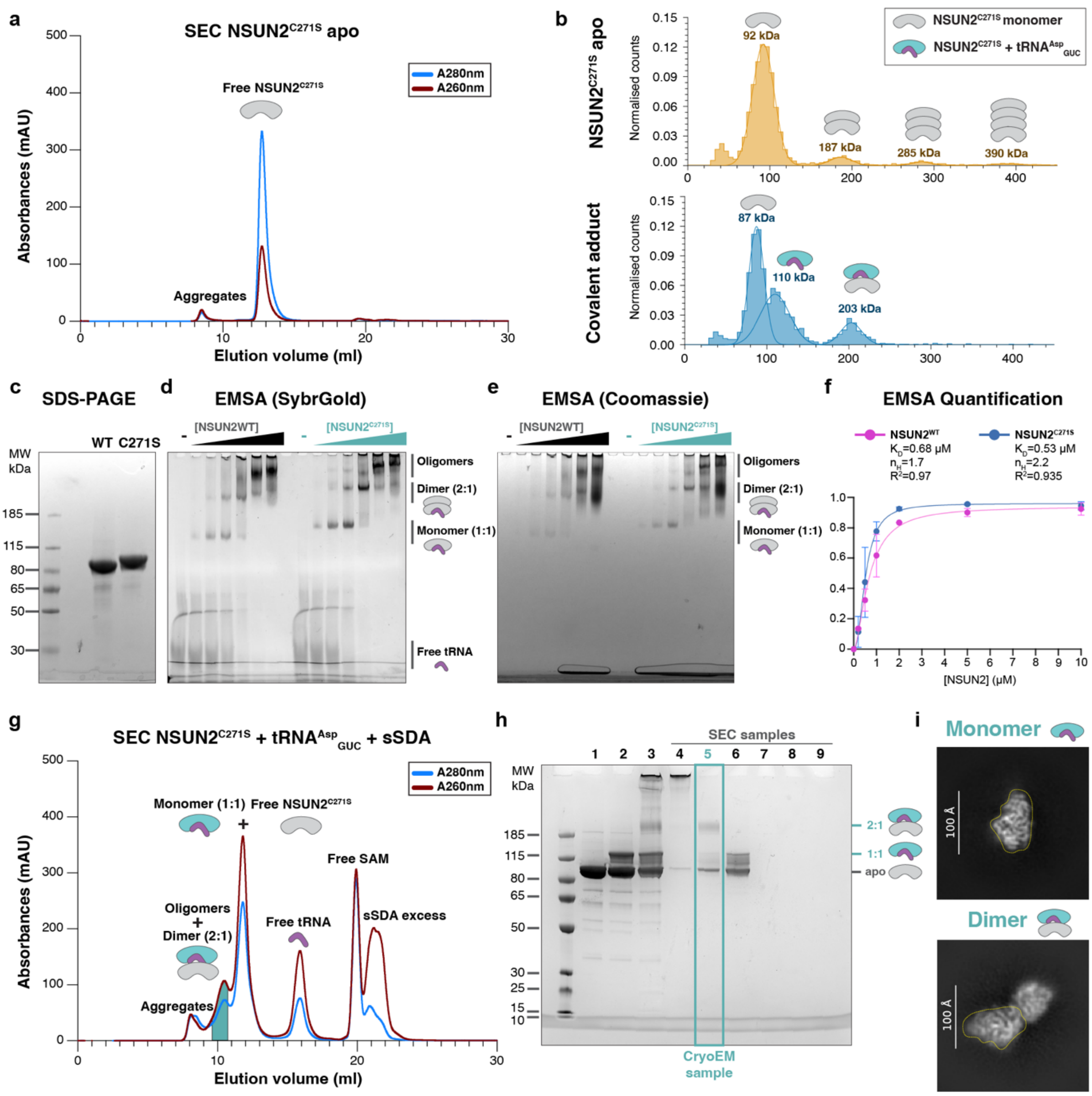

**Extended Data Fig 2 | Purification of an active NSUN2-tRNA$^{Asp}$$_{GUC}$ complex in a post-catalytic state.**

**a**, Size-exclusion chromatography (SEC) profile of apo NSUN2$^{C271S}$ showing a predominant monomeric peak and a minor aggregate fraction (UV absorbance at 280 nm and 260 nm).

**b**, Mass photometry of apo NSUN2$^{C271S}$ (top) and the covalent NSUN2$^{C271S}$-tRNA$^{Asp}$$_{GUC}$ adduct formed in the presence of SAM (bottom). The apo sample shows a predominant monomeric species and additional oligomeric populations with molecular masses consistent with 2, 3, and 4 copies of NSUN2. Upon tRNA adduct formation, species corresponding to monomeric NSUN2, a 1:1 NSUN2-tRNA complex, and a 2:1 NSUN2:tRNA complex are observed.

**c**, Coomassie-stained SDS–PAGE analysis of purified wild-type NSUN2$^{WT}$ and NSUN2$^{C271S}$, showing a difference in molecular weight due to two different tagging strategies.

**d,e,** Electrophoretic mobility shift assays (EMSA) comparing binding of NSUN2$^{WT}$ and NSUN2$^{C271S}$ to tRNA$^{Asp}$$_{GUC}$, visualised by SybrGold staining (**d**) and Coomassie staining (**e**), showing formation of monomeric and higher-order NSUN2–tRNA species at increasing protein concentrations. The RNA sequences are summarized in Supplementary Table S3.

**f,** Quantification of EMSA binding (from **d,e**). Quantification was performed on two independent experiments (n = 2).

**g,** SEC profile of the NSUN2$^{C271S}$-tRNA$^{Asp}$$_{GUC}$ reconstitution reaction after mild stabilisation with sulfo-SDA (sSDA), showing fractions corresponding to monomeric NSUN2$^{C271S}$, 1:1 complex, dimeric species (2:1 NSUN2:tRNA) and higher-order oligomers, as well as free tRNA, SAM, and excess crosslinker.

**h,** SDS–PAGE analysis of SEC fractions from **g**, with the fraction used for cryo-EM grid preparation indicated.

**i,** Representative 2D class averages illustrating monomeric and dimeric particles observed for the NSUN2-tRNA$^{Asp}$$_{GUC}$ preparation.

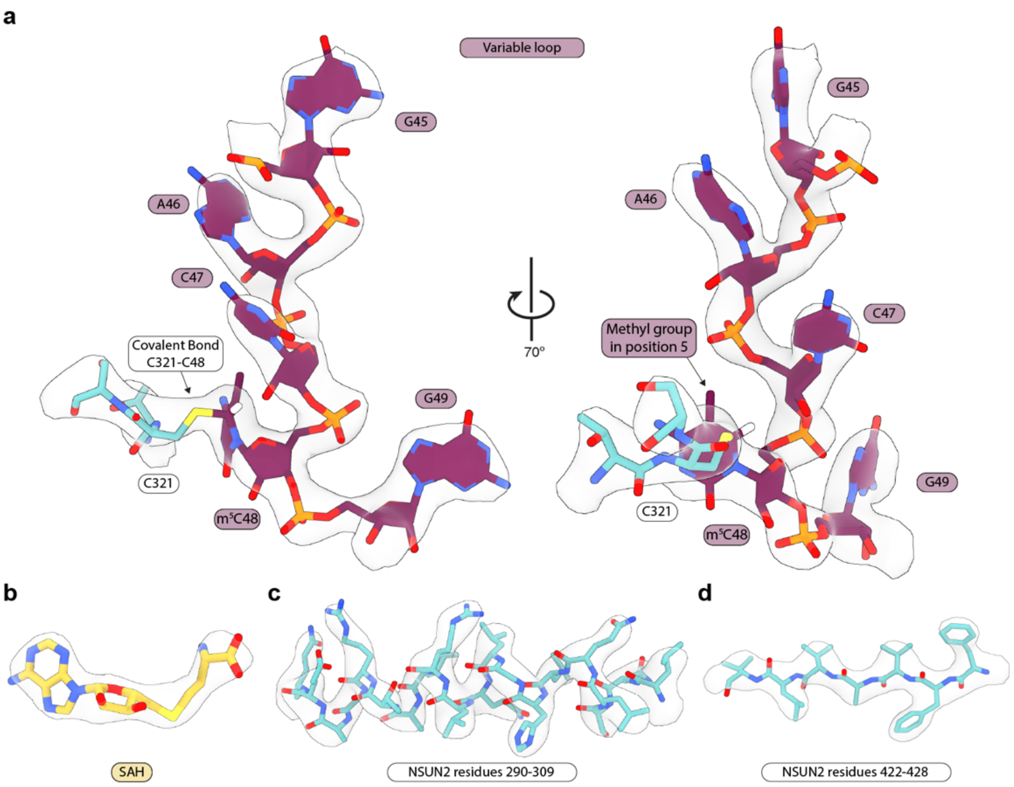

**Extended Data Fig. 3 | Map features of the post-catalytic active site.**

**a,** Cryo-EM density for the tRNA variable-loop in the NSUN2 catalytic centre, showing m$^5$C48 covalently linked to Cys321 (map shown as transparent grey).

**b,** Cryo-EM density for S-adenosylhomocysteine (SAH) in the cofactor-binding pocket.

**c,d,** Representative regions of the final reconstruction illustrating map quality and fit of the model into the map (map shown as transparent grey).

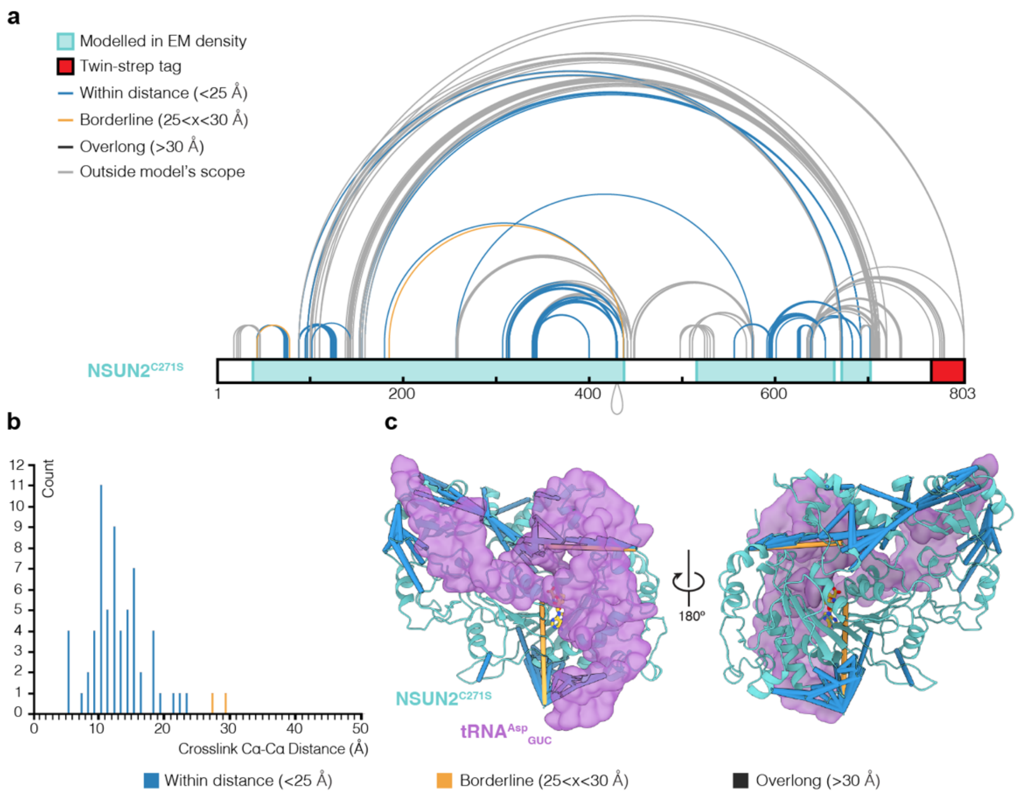

**Extended Data Fig.4 | Protein-protein chemical crosslinking mass spectrometry analysis supports and assists the EM model building of the NSUN2-tRNA complex.**

**a,** Chemical crosslink map of sulfoSDA-derived crosslinks mapped onto the NSUN2$^{C271S}$ sequence.

**b,** Distance distribution of Cα–Cα distances for the 64 residue pairs mapping to the NSUN2-tRNA complex structure. A total of 62 crosslinks (99%) fall within the sulfoSDA distance restraints (<25 Å), confirming the model's accuracy. Histogram bars are coloured by distance.

**c,** sulfoSDA-crosslinks mapped to the NSUN2-tRNA model, coloured by distance. NSUN2$^{C271S}$ is shown in teal, and the tRNA$^{Asp}{}_{GUC}$ in purple.

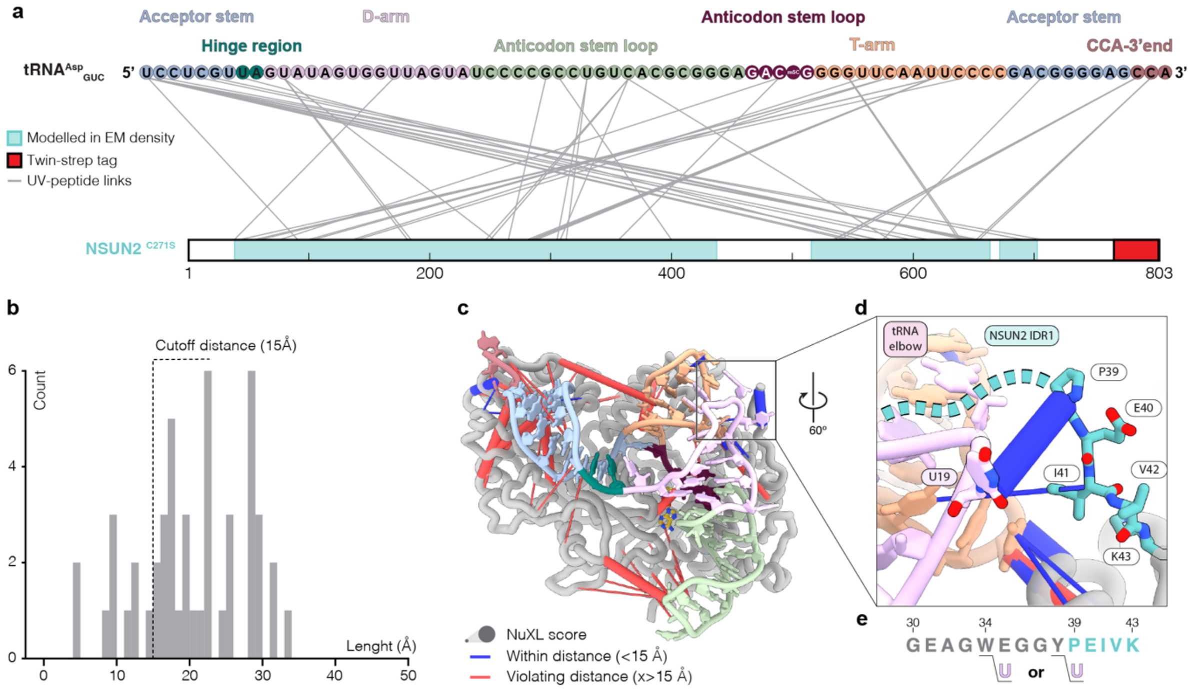

**Extended Data Fig.5 | Protein-RNA UV crosslinking mass spectrometry analysis supports and assists the EM model building of the NSUN2-tRNA complex.**

**a,** Barplot of UV-derived crosslinks mapped onto the NSUN2$^{C271S}$ and tRNA$^{Asp}_{GUC}$ sequence. The links, coloured in gray, are mapped according to the shortest possible distance between the protein peptide and the corresponding nucleotide adduct.

**b,** Histogram of Protein–RNA distances for the 45 modelled crosslinks. Distances are calculated between the protein Cα and the closest nucleobase ring atom. The cutoff for the zero-length UV-induced link was set to 15 Å.

**c,** Overall representation of UV crosslinks mapped to the NSUN2–tRNA model, coloured by distance and with thickness proportional to the corresponding NuXL score. NSUN2$^{C271S}$ is shown in grey, and the tRNA$^{Asp}_{GUC}$ is coloured by domains. The anticodon stem is coloured in green, the variable loop in burgundy, the D-arm in pink, the T-arm in orange, the hinge region in dark turquoise, the acceptor stem in light blue, and the CCA in coral.

**d,** Close-up representation of UV crosslinks mapped to the NSUN2–tRNA model, coloured by distance and with thickness proportional to the corresponding NuXL score. NSUN2$^{C271S}$ is shown in grey, and the tRNA$^{Asp}_{GUC}$ is coloured by domains. The T-arm is coloured orange, the D-arm pink, and the NSUN2$^{C271S}$ IDR1 teal.

**e,** Peptide-Nucleobase adducts detected with MS for the link highlighted in panel **d** reveal link formation between W34 and Y38 and a uracil. These protein residues are part of an unmodelled portion of NSUN2$^{C271S}$ IDR1. Modelled amino acids are coloured in teal, the crosslinked T-arm uracil is pink, and the unmodelled amino acids in gray.

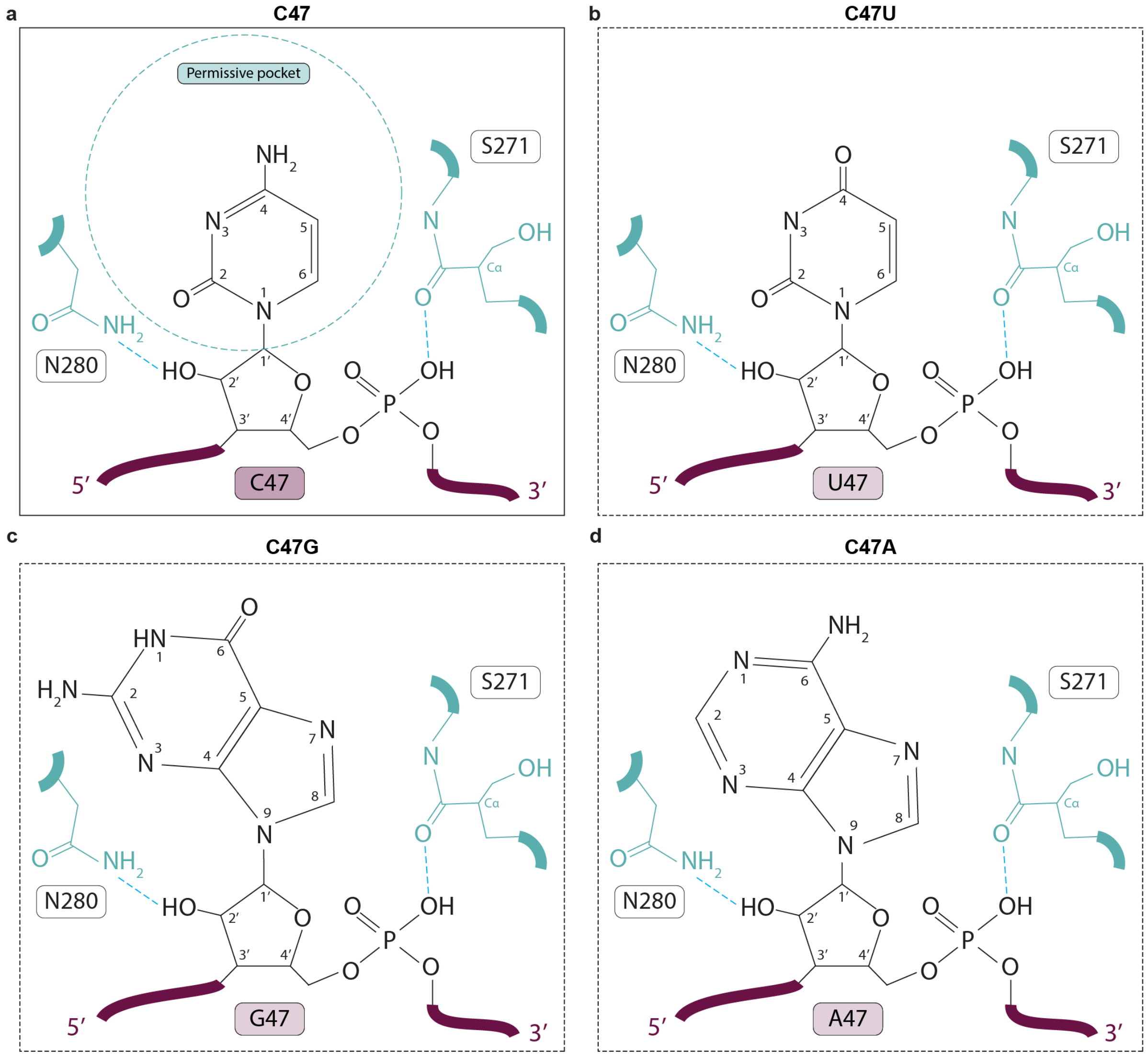

**Extended Data Fig.6 | Position 47 (−1 relative to the $m^5C48$ site) forms a permissive pocket accommodating any nucleotide.**

**a,** C47 is stabilized by backbone-mediated hydrogen bonds with $NSUN2^{C271S}$ (coloured in teal), involving Asn280 and the backbone carbonyl of Ser271 (Cys271 in the wild type), consistent with the absence of nucleotide specificity at this position.

**b-d,** Structural views illustrating how different nucleotides at position 47 are accommodated within the permissive pocket, with backbone-mediated interactions supporting nucleotide-independent recognition.

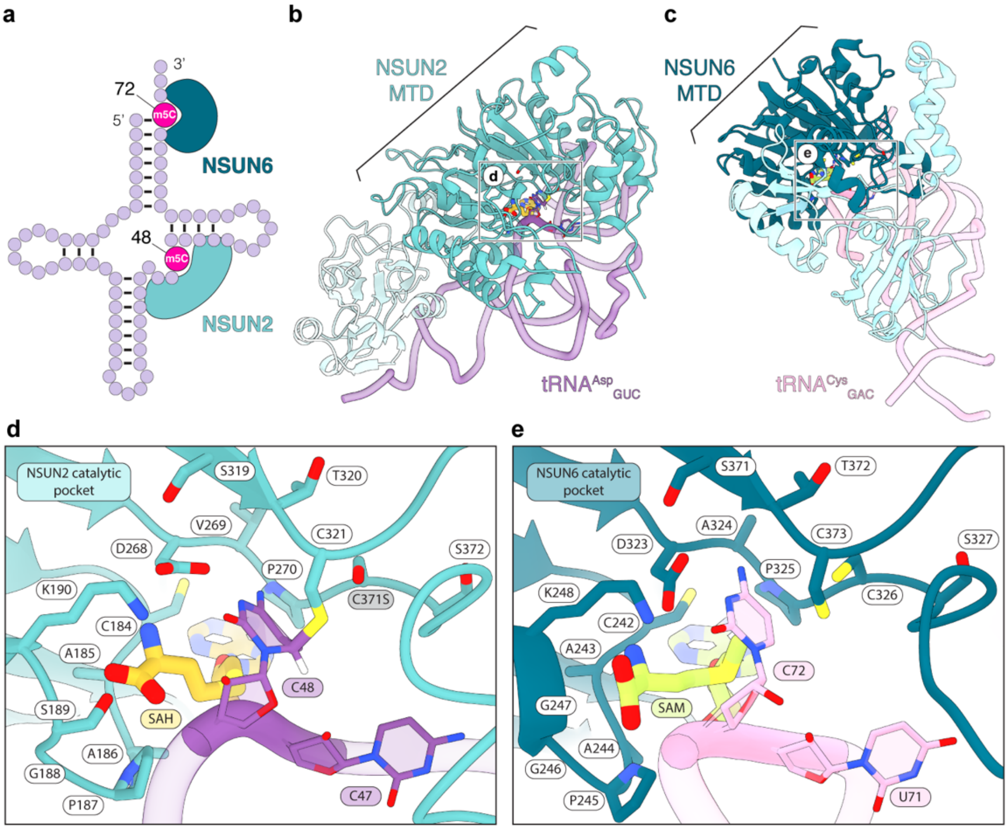

**Extended Data Fig.7 | NSUN2-NSUN6 comparison.**

**a,** Schematic comparison of NSUN2 and NSUN6 binding modes on tRNA. NSUN2 methylates position 48 within the variable loop, whereas NSUN6 methylates position 72 within the acceptor stem.

**b,** Cartoon representation of NSUN2$^{C271S}$ binding to tRNA$^{Asp}_{GUC}$ showing the orientation corresponding to panel **d**. NSUN2$^{C271S}$ is shown in teal, and the tRNA in purple.

**c,** Cartoon representation of NSUN6 binding to tRNA$^{Cys}_{GAC}$ (PDB: 5WWS) showing the orientation corresponding to panel **e**. NSUN6 is shown in dark teal, and the tRNA in pink.

**d,e,** Structural comparison of NSUN2 and NSUN6 catalytic centers. Conserved residues are annotated. **d,** NSUN2$^{C271S}$ catalytic center bound to tRNA$^{Asp}_{GUC}$ and **e,** NSUN6 catalytic center bound to the tRNA$^{Cys}_{GAC}$ (PDB: 5WWS). NSUN2$^{C271S}$ is shown in teal; NSUN6 dark teal; the tRNA$^{Asp}_{GUC}$ purple, the tRNA$^{Cys}_{GAC}$ pink; SAH in gold, and SAM in lime.

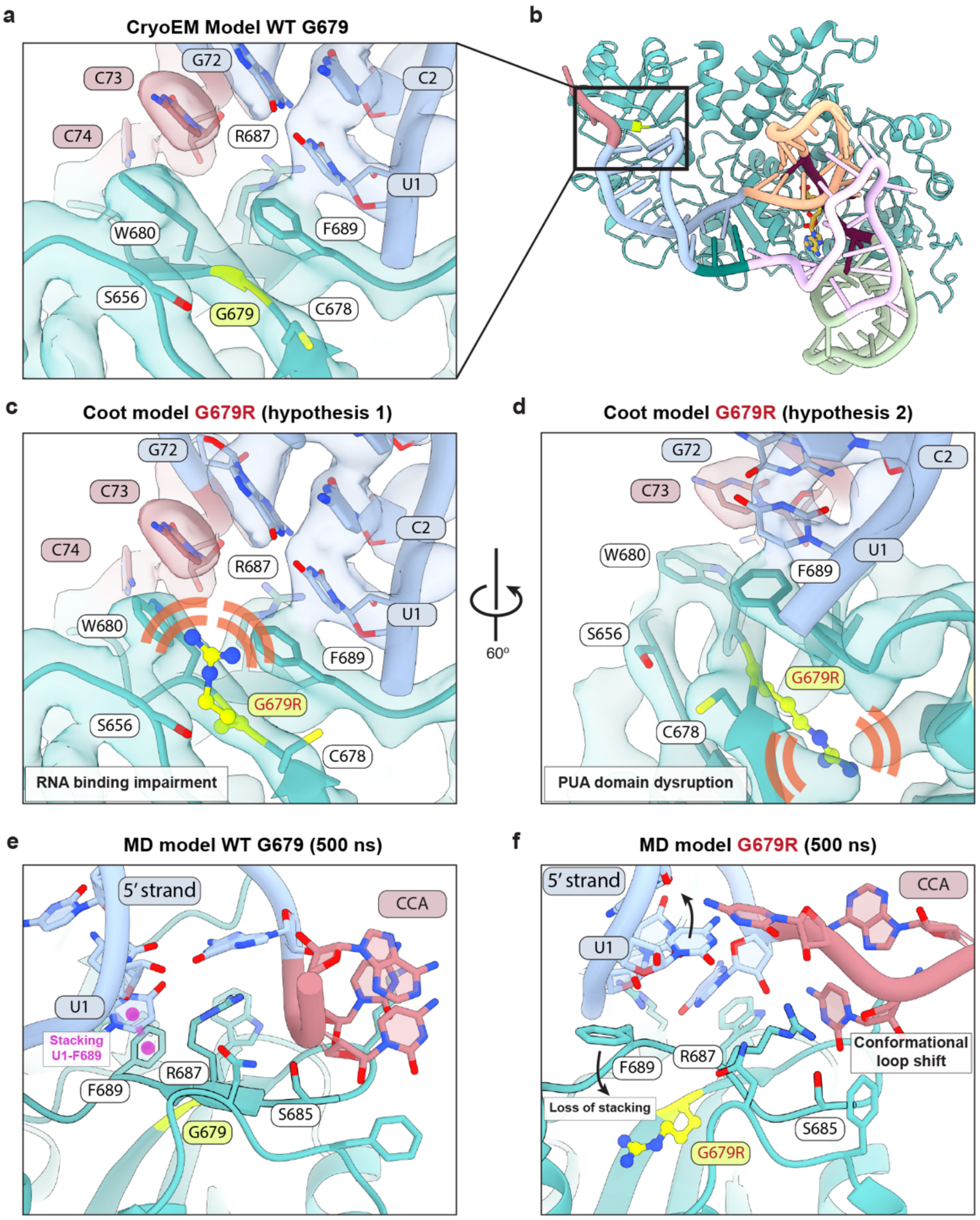

**Extended Data Fig. 8 | Impact of Gly679Arg (G679R) mutation on tRNA binding.**
**a,** Zoom-in view of the PUA domain highlighting G679, located within a β-strand adjacent to the tRNA acceptor stem. The U1–G72 base pair is stabilised by stacking interactions with aromatic residues (Phe689 and Trp680) and hydrogen bonding with Arg687. Gly679 is shown in fluorescent yellow.

**b,** Overall view of Gly679 within the cryo-EM structure. NSUN2$^{C271S}$ is shown in teal, and the tRNA is coloured by structural domains as indicated.

**c,** Manual modelling in Coot of the G679R mutation within the PUA domain. In the first hypothesis, the arginine side chain is oriented toward the tRNA-binding interface, where it would sterically hinder proper positioning of the acceptor stem and impair RNA binding.

**d,** Manual modelling in Coot of the G679R mutation within the PUA domain. In the second hypothesis, the arginine side chain is oriented toward the β-sheet core of the PUA domain (β-strand numbering indicated), where it could disrupt β-sheet packing and domain folding, thereby affecting accommodation of the tRNA acceptor stem and CCA terminus.

**e,f,** Representative structures from 500-ns molecular dynamics simulations of the wild-type (**e**) and G679R mutant (**f**). Structurally, replacement of glycine with arginine introduces a bulky, positively charged guanidinium side chain into a tightly packed region of the PUA domain. The increased steric demand likely disrupts local packing interactions and imposes conformational strain, while the altered electrostatic environment may perturb neighbouring aromatic and RNA-contacting residues. The G679R substitution induces displacement of the loop comprising residues 685–690, disrupting the stacking interaction between U1 and Phe689 observed in the wild-type structure. This rearrangement provides a structural basis for reduced tRNA binding in the mutant. Collectively, these effects enhance backbone mobility within the RNA-binding domain, weaken optimal protein-RNA interactions, and contribute to moderate destabilization of the complex. Overall, the data indicate that the G679R mutation moderately destabilizes the NSUN2-tRNA complex, potentially influencing protein-RNA interaction dynamics.

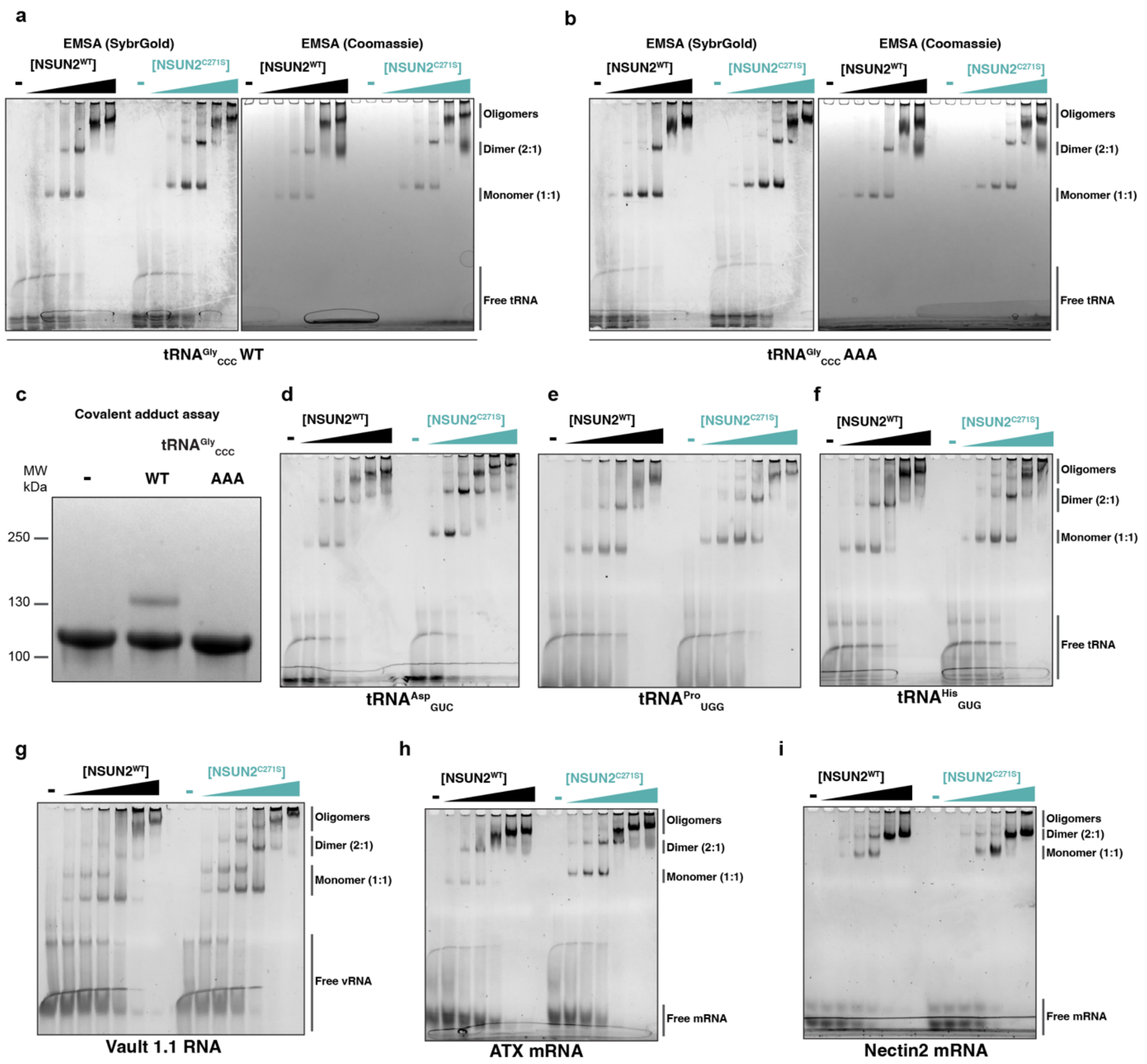

**Extended Data Fig. 9 | EMSA analysis of various NSUN2 RNA substrates.**

**a,** Electrophoretic mobility shift assay (EMSA) stained with SybrGold or Coomassie, assessing binding of NSUN2$^{WT}$ (black) and NSUN2$^{C271S}$ (teal) to a full-length wild-type tRNA$^{Gly}_{CCC}$.

**b,** Electrophoretic mobility shift assay (EMSA) stained with SybrGold or Coomassie, assessing binding of NSUN2$^{WT}$ (black) and NSUN2$^{C271S}$ (teal) to a full-length mutated tRNA$^{Gly}_{CCC}$ (AAA instead of CCC at positions 46-47-48 within the variable loop).

**c,** Covalent adduct formation assay using NSUN2$^{C271S}$ mutant and stained with Coomassie showing that the full-length mutated tRNA$^{Gly}_{CCC}$ substrate (AAA instead of CCC at positions 46-47-48 within the variable loop) does not support productive covalent complex formation under the conditions tested, in contrast to full-length wild-type tRNA$^{Gly}_{CCC}$.

**d-i,** Electrophoretic mobility shift assay (EMSA) stained with SybrGold assessing binding of $NSUN2^{WT}$ (black) and $NSUN2^{C271S}$ (teal) to a **d**, wild-type $tRNA^{Asp}_{GUC}$, **e**, wild-type $tRNA^{Pro}_{UGG}$, **f**, wild-type $tRNA^{His}_{GUG}$, **g**, wild-type vault 1.1 RNA, **h**, wild-type ATX mRNA, and **i**, Nectin2 mRNA. The RNA sequences are summarized in Supplementary Table S3.

**SUPPLEMENTARY TABLES (with legends)**

| | pXL-MS $NSUN2^{C271S}$-$tRNA^{Asp}_{GUC}$ | Non-crosslinked $NSUN2^{C271S}$-$tRNA^{Asp}_{GUC}$ |
|---|---|---|
| **Data collection and processing** | | |
| Magnification | 215’000 | 215’000 |
| Voltage (kV) | 300 | 300 |
| Electron exposure (e–/Å$^2$) | 60 | 60 |
| Defocus range (µm) | 0.8-1.8 | 0.5-1.5 |
| Pixel size (Å) | 0.582 | 0.582 |
| Symmetry imposed | C1 | C1 |
| Initial particle images (no.) | 762’007 | 2’133’046 |
| Final particle images (no.) | 33’531 | 182’859 |
| Map resolution (Å) | 3.1 | 3.0 |
| FSC threshold | 0.143 | 0.143 |
| Map resolution range (Å) | 2.8/6.2 | 2.6/6.2 |
| **Refinement** | | |
| Initial model used (PDB code) | | |
| Model resolution (Å) | 3.3 | |
| FSC threshold | 0.5 | |
| Model resolution range (Å) | | |
| Map sharpening *B* factor (Å$^2$) | -15 | |
| Model composition | | |
| Non-hydrogen atoms | 6306 | |
| Protein residues | 579 | |
| Ligands | 1 | |
| RNA/DNA nucleotides | 75 | |
| *B* factors (Å$^2$) | | |
| Protein | 98.94 | |
| Ligand | | |
| R.m.s. deviations | | |

| | |
|---|---|
| Bond lengths (Å) | 0.007 |
| Bond angles (°) | 0.595 |
| Validation | |
| MolProbity score | 1.90 |
| Clashscore | 12.66 |
| Poor rotamers (%) | 0.38 |
| Ramachandran plot | |
| Favored (%) | 95.80 |
| Allowed (%) | 4.02 |
| Disallowed (%) | 0.17 |

**Table S1: Cryo-EM data collection, refinement, and validation statistics**

| Primer number | Primer name | Sequence |
|---|---|---|
| **1** | NSUN2-3CSII_C271S_fwd | GCGACGTTCCGAGCTCT GGCGATGGCACC |
| **2** | NSUN2-3CSII_C271S_rvs | GGTGCCATCGCCAGAG CTCGGAACGTCGC |

**Table S2: List of primers used to generate NSUN2$^{C271S}$ mutagenesis.**

| Name | Sequence (5'-3') | Gene Locus |
|---|---|---|
| **$tRNA^{Asp}_{GUC}$** | TAATACGACTCACTATAGGGTCCTC GTTAGTATAGTGGTTAGTATCCCCG CCTGTCACGCGGGAGACCGGGGTTC AATTCCCCGACGGGGAGCCA | tRNA-Asp-GUC-1-1 |
| **T-arm + variable loop $tRNA^{Asp}_{GUC}$** | TAATACGACTCACTATAGGGACCGG GGTTCAATTCCCCG | tRNA-Asp-GUC-1-1 |
| **$tRNA^{Pro}_{UGG}$** | TAATACGACTCACTATAGGGGGCTC GTTGGTCTAGGGGTATGATTCTCGC TTTGGGTGCGAGAGGTCCCGGGTTC AAATCCCGGACGAGCCCCCA | tRNA-Pro-UGG-3-5 |
| **$tRNA^{Gly}_{CCC}$ WT** | TAATACGACTCACTATAGGGGCGCC GCTGGTGTAGTGGTATCATGCAAGA TTCCCATTCTTGCGACCCGGGTTCGA TTCCCGGGCGGCGCACCA | tRNA-Gly-CCC-2-1 |
| **$tRNA^{Gly}_{CCC}$ AAA** | TAATACGACTCACTATAGGGGCGCC GCTGGTGTAGTGGTATCATGCAAGA TTCCCATTCTTGCGAAAAGGGTTCG ATTCCCGGGCGGCGCACCA | tRNA-Gly-CCC-2-1 |
| **$tRNA^{His}_{GUG}$** | TAATACGACTCACTATAGGGCCGTG ATCGTATAGTGGTTAGTACTCTGCG TTGTGGCCGCAGCAACCTCGGTTCG AATCCGAGTCACGGCACCA | tRNA-His-GUG-1-7 |
| **vtRNA 1.1** | TAATACGACTCACTATAGGGGGGCT GGCTTTAGCTCAGCGGTTACTTCGA CAGTTCTTTAATTGAAACAAGCAAC CTGTCTGGGTTGTTCGAGACCCGCG GGCGCTCTCCAGTCCTTTT | vtRNA 1.1 |
| **ATX mRNA** | TAATACGACTCACTATAGGGATATT TTTATATTGTTTTTGTATTTATTAATT TGAAACCAGGACATTAAAAATG | chr8: 119557427-119557480 |

| | | |
|---|---|---|
| **NECTIN2 mRNA** | TAATACGACTCACTATAGGGCCCTC TGCATCTCCAAAGAGGGC | chr19: 44871916-44871938 |

**Table S3: List of RNA constructs generated by T7 *in vitro* transcription.**

tRNA's sequences have been selected from the genomic tRNA database: https://gtrnadb.org/

| ## | Structure 1 | Distance (Å) | Structure 2 |
|---|---|---|---|
| **1** | B:A 46[ N6 ] | 3.26257 | A:ASN 291[ OD1] |
| **2** | B:A 46[ N6 ] | 3.01906 | A:GLN 294[ OE1] |
| **3** | B:U 58[ N3 ] | 3.80372 | A:ASP 282[ OD1] |
| **4** | B:A 65[ N3 ] | 3.60949 | A:GLU 517[ OE2] |
| **5** | B:G 51[ OP2] | 2.94514 | A:ARG 133[ NE ] |
| **6** | B:G 50[ OP2] | 3.38468 | A:ARG 133[ NH1] |
| **7** | B:G 51[ OP2] | 2.86507 | A:ARG 137[ NH1] |
| **8** | B:G 52[ OP2] | 3.54078 | A:ARG 137[ NH2] |
| **9** | B:U 54[ O4 ] | 3.71258 | A:LYS 138[ NZ ] |
| **10** | B:G 51[ OP1] | 3.02565 | A:HIS 146[ NE2] |
| **11** | B:G 50[ O3'] | 3.62755 | A:ARG 160[ NH1] |
| **12** | B:G 51[ OP1] | 3.3794 | A:ARG 160[ NH1] |
| **13** | B:G 49[ OP1] | 2.99909 | A:GLU 162[ N ] |
| **14** | B:G 49[ OP2] | 3.75263 | A:SER 165[ OG ] |
| **15** | B:A 9[ O2'] | 3.87137 | A:ASN 218[ ND2] |
| **16** | B:C 66[ OP1] | 2.80109 | A:LYS 229[ NZ ] |
| **17** | B:C 47[ OP2] | 3.61173 | A:SER 271[ N ] |
| **18** | B:C 47[ O2'] | 3.63674 | A:ASN 280[ ND2] |
| **19** | B:G 45[ OP1] | 2.99399 | A:LYS 286[ NZ ] |
| **20** | B:G 43[ O2'] | 3.57116 | A:GLN 294[ NE2] |
| **21** | B:C 29[ O3'] | 3.83033 | A:ARG 301[ NH1] |
| **22** | B:G 30[ OP2] | 2.90219 | A:ARG 301[ NH1] |
| **23** | B:C 29[ OP1] | 3.18405 | A:ARG 301[ NH2] |
| **24** | B:G 43[ OP1] | 3.50034 | A:LYS 369[ NZ ] |
| **25** | B:C 66[ O2'] | 3.60538 | A:ARG 551[ NH2] |
| **26** | B:G 68[ OP2] | 2.92123 | A:ARG 558[ NE ] |
| **27** | B:G 69[ OP2] | 3.81469 | A:ARG 558[ NH1] |
| **28** | B:G 68[ OP2] | 2.96333 | A:ARG 558[ NH2] |
| **29** | B:G 67[ OP1] | 3.80965 | A:GLN 559[ NE2] |
| **30** | B:G 68[ OP2] | 3.58186 | A:GLN 559[ NE2] |
| **31** | B:G 67[ OP2] | 2.59768 | A:TYR 561[ OH ] |

| | | | |
|---|---|---|---|
| **32** | B:C 66[ O5'] | 3.8936 | A:ASN 582[ ND2] |
| **33** | B:A 65[ O2'] | 3.58639 | A:ASN 582[ ND2] |
| **34** | B:U 1[ O4 ] | 3.52964 | A:ARG 602[ NH2] |
| **35** | B:C 74[ O2 ] | 2.80433 | A:ALA 684[ N ] |
| **36** | B:C 74[ O2 ] | 3.66576 | A:SER 685[ N ] |
| **37** | B:C 74[ O2 ] | 3.43048 | A:SER 685[ OG ] |
| **38** | B:U 1[ O2 ] | 3.89989 | A:ARG 687[ NH2] |
| **39** | B:5MC 48[ N4 ] | 2.82173 | A:ASP 268[ OD1] |
| **40** | B:5MC 48[ N4 ] | 2.61435 | A:VAL 269[ O ] |
| **41** | B:5MC 48[ O3'] | 3.33355 | A:LYS 279[ NZ ] |
| **42** | B:5MC 48[ O2 ] | 3.71718 | A:LYS 190[ NZ ] |
| **43** | B:5MC 48[ OP3] | 2.77641 | A:ARG 220[ NH2] |

**Table S4: Hydrogen bonds listing coming from PEDBePISA analysis of NSUN2$^{C271S}$ – tRNA$^{Asp}{}_{GUC}$ surface of interaction. (https://www.ebi.ac.uk/pdbe/pisa/) (No disulfide bonds, no salt bridges were found)**

| ## | Structure 1 | Distance (Å) | Structure 2 |
|---|---|---|---|
| **1** | B:5MC 48[ C6 ] | 1.87 | A:CYS 321[ SG ] |

**Table S5: Covalent bonds listing coming from PEDBePISA analysis of NSUN2$^{C271S}$ – tRNA$^{Asp}_{GUC}$ surface of interaction. (https://www.ebi.ac.uk/pdbe/pisa/) (No disulfide bonds, no salt bridges were found)**

| PDB Structure | RMSD | | RMSF | |
|---|---|---|---|---|
| | **Protein backbone** | **RNA** | **Protein backbone** | **RNA** |
| NSUN2-tRNA | 2.54 ± 0.43 | 5.50 ± 1.03 | 1.24 ± 0.14 | 2.84 ± 0.37 |
| NSUN2-tRNA (G679R) | 2.99 ± 0.33 | 4.97 ± 0.58 | 1.28 ± 0.05 | 3.19 ± 0.53 |

**Table S6: RMSD (Root Mean Square Deviation) and RMSF (Root Mean Square Fluctuation) values of protein and RNA in NSUN2-tRNA complex and its G679R mutant.** The calculations are based on the backbone carbon atoms of protein and the phosphate atoms of tRNA. The values represent the average RMSD and RMSF obtained from two independent simulations.

#### SUPPLEMENTARY DATA (with legends)

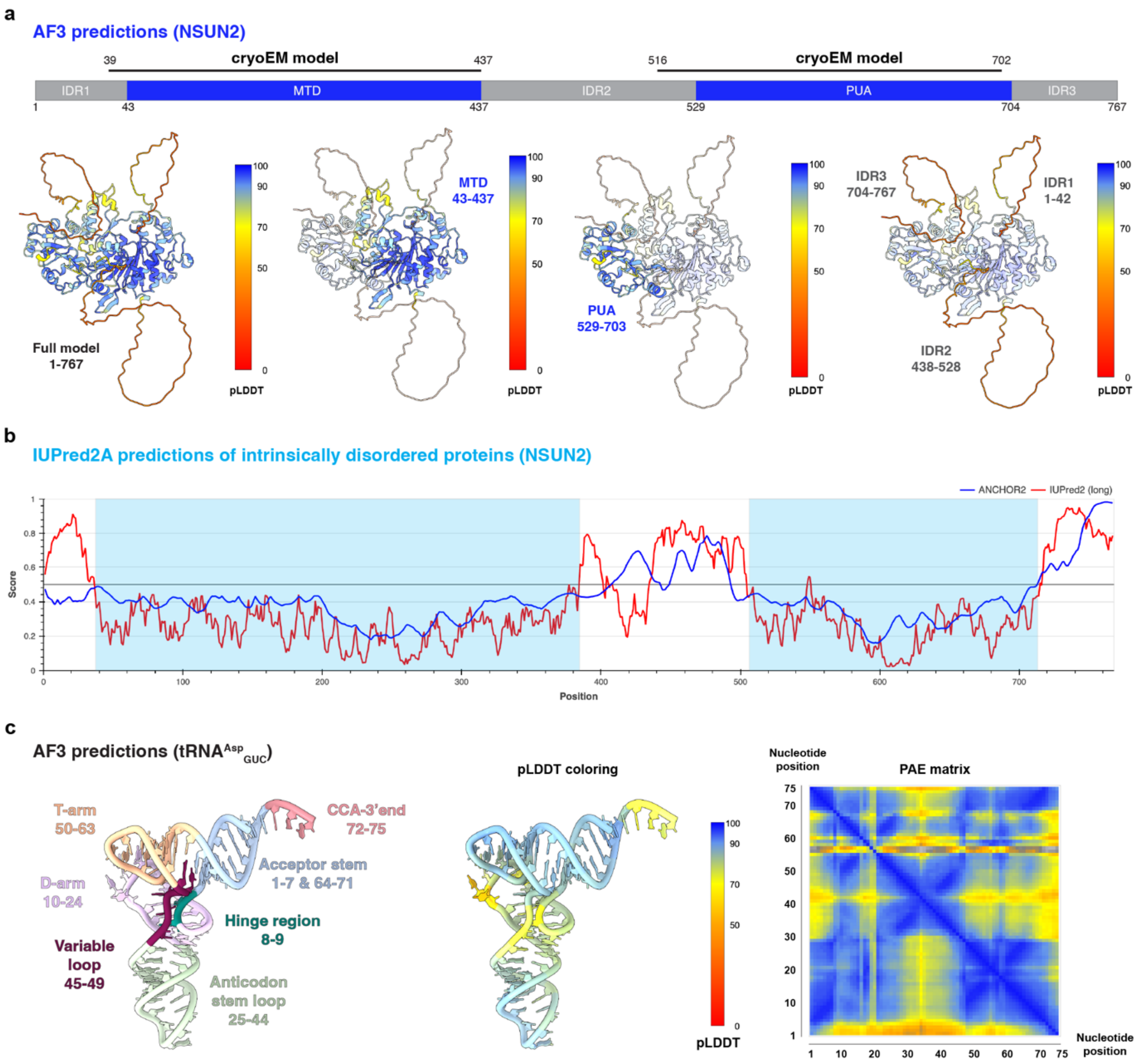

**Supplementary Data Fig.1 | Structural predictions of NSUN2 and the tRNA$^{Asp}{}_{GUC}$.**

**a,** AlphaFold3 structure predictions of NSUN2$^{WT}$ highlighting the folded methyltransferase domain (MTD) and PUA domain, as well as three intrinsically disordered regions (IDRs), consistent with the regions resolved in the cryoEM model. Panels illustrate the domain organisation and folding by selective highlighting of individual regions.

**b,** IUPred2A disorder prediction of NSUN2$^{WT}$ showing folded MTD and PUA domains highlighted in teal and three IDRs, consistent with AlphaFold3 predictions and our cryoEM model.

**c,** AlphaFold3 prediction of human $tRNA^{Asp}_{GUC}$ highlighting the canonical secondary structure elements, including the acceptor stem, D-arm, anticodon loop, variable loop, T-arm, and CCA 3′ end. The same model coloured by per-residue confidence (pLDDT). Predicted aligned error (PAE) map indicating overall high confidence in the relative positioning of tRNA domains, consistent with a well-defined global fold.

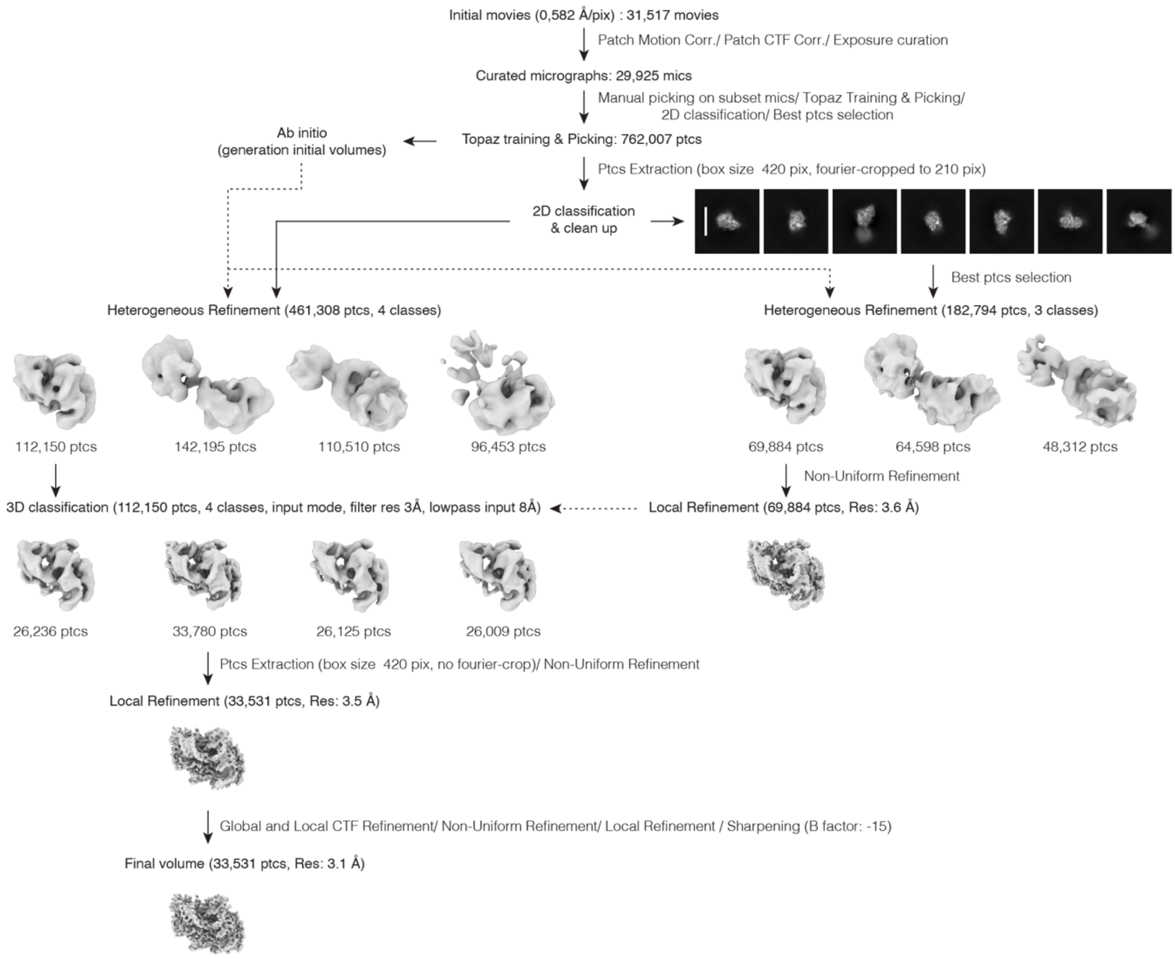

**Supplementary Data Fig.2 | CryoEM workflow of the pXL-EM sample.**

Movie stacks were imported, motion-corrected, and CTF-estimated prior to manual micrograph curation. Topaz particle picking was trained on a manually curated subset of particles to generate an initial particle pool, which was subjected to 2D classification. After a second round of Topaz training and picking, extracted particles were used for Ab-initio reconstruction followed by Heterogeneous Refinement to obtain initial volumes corresponding to different oligomeric states of the complex. The reconstruction corresponding to the 1:1 stoichiometry was further refined using Non-Uniform and Local Refinement, yielding a map at 3.6 Å resolution. In parallel, the full Topaz-picked particle set was processed by direct heterogeneous refinement to expand the particle pool. Particles assigned to the 1:1 reconstruction were selected and subjected to 3D classification using the 3.6 Å map as a reference. The best 3D class (33,531 particles) was selected for final refinement and post-processing, resulting in a 3.1 Å resolution map (FSC 0.143). Dashed arrows indicate the propagation of volumes between steps, whereas solid arrows represent particle sets.

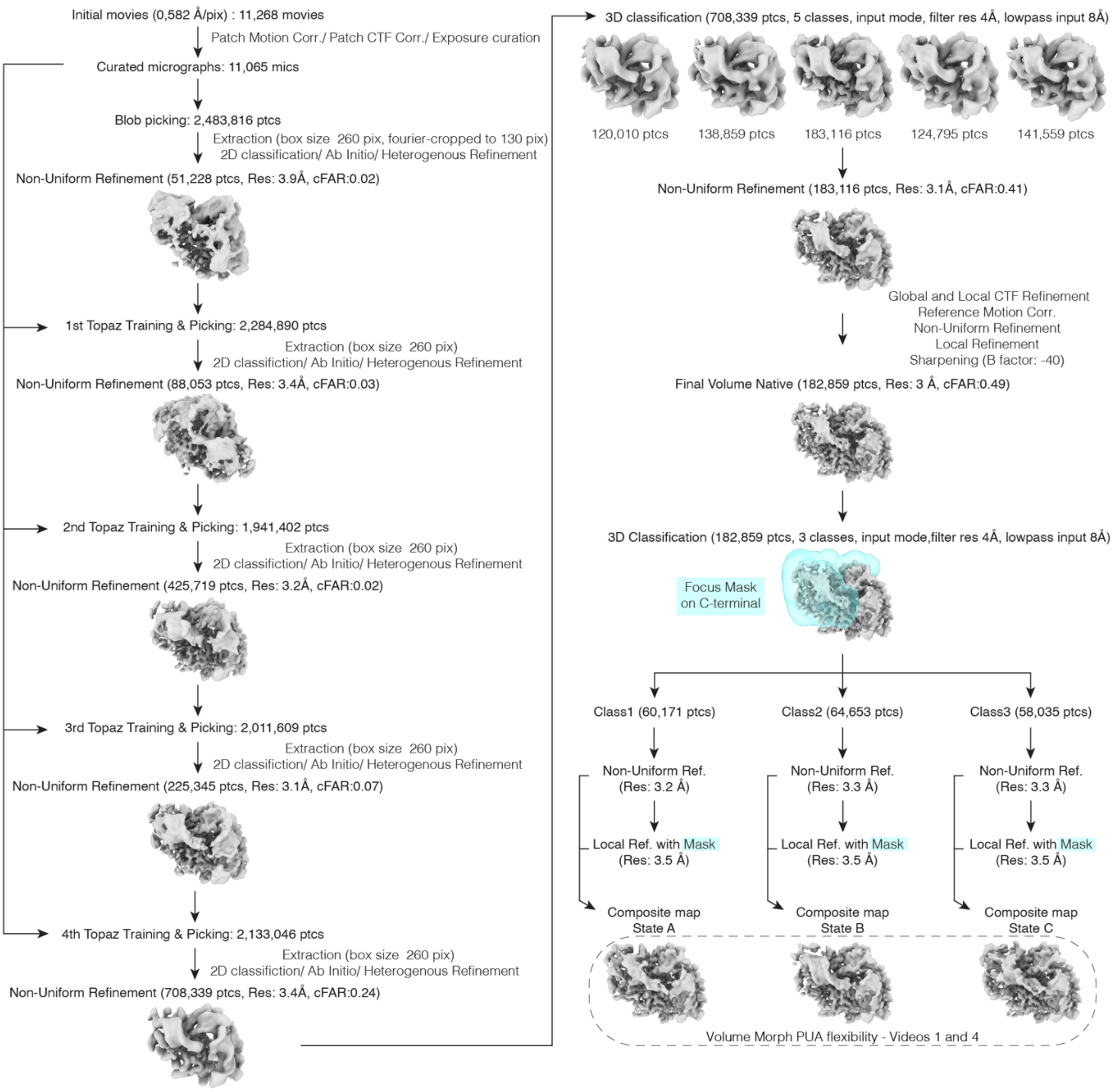

**Supplementary Data Fig.3 | CryoEM workflow of the non-crosslinked sample.**
Movie stacks were imported, motion-corrected, and CTF-estimated prior to manual micrograph curation. An initial particle set obtained by blob picking was cleaned through iterative 2D classification and used to train a TOPAZ neural network picker. Four cycles of TOPAZ picking, 2D classification, Ab-Initio reconstruction, Heterogeneous Refinement, and Non-Uniform Refinement were performed to progressively improve map quality. The final TOPAZ picking yielded ~2.1 million particles, which were subjected to broad Heterogeneous Refinement to remove contaminants and poorly resolved classes. Selected particles were refined by Non-Uniform Refinement, followed by 3D classification to isolate the most homogeneous subset. The best 3D class (183,859 particles) was selected for final refinement and post-processing, resulting in a final reconstruction at 3.0 Å resolution (FSC 0.143). To

visualize the conformational flexibility of the PUA domain within the non-crosslinked complex, the final reconstruction was subjected to an additional round of 3D classification using a focused mask encompassing the C-terminal region of the complex (the PUA domain and the tRNA acceptor stem) to further resolve structural heterogeneity. Each resulting class was subsequently refined using both Non-Uniform Refinement with the solvent mask and Local Refinement with the focused mask. The three volumes obtained from this procedure were then used to generate the morph in ChimeraX.

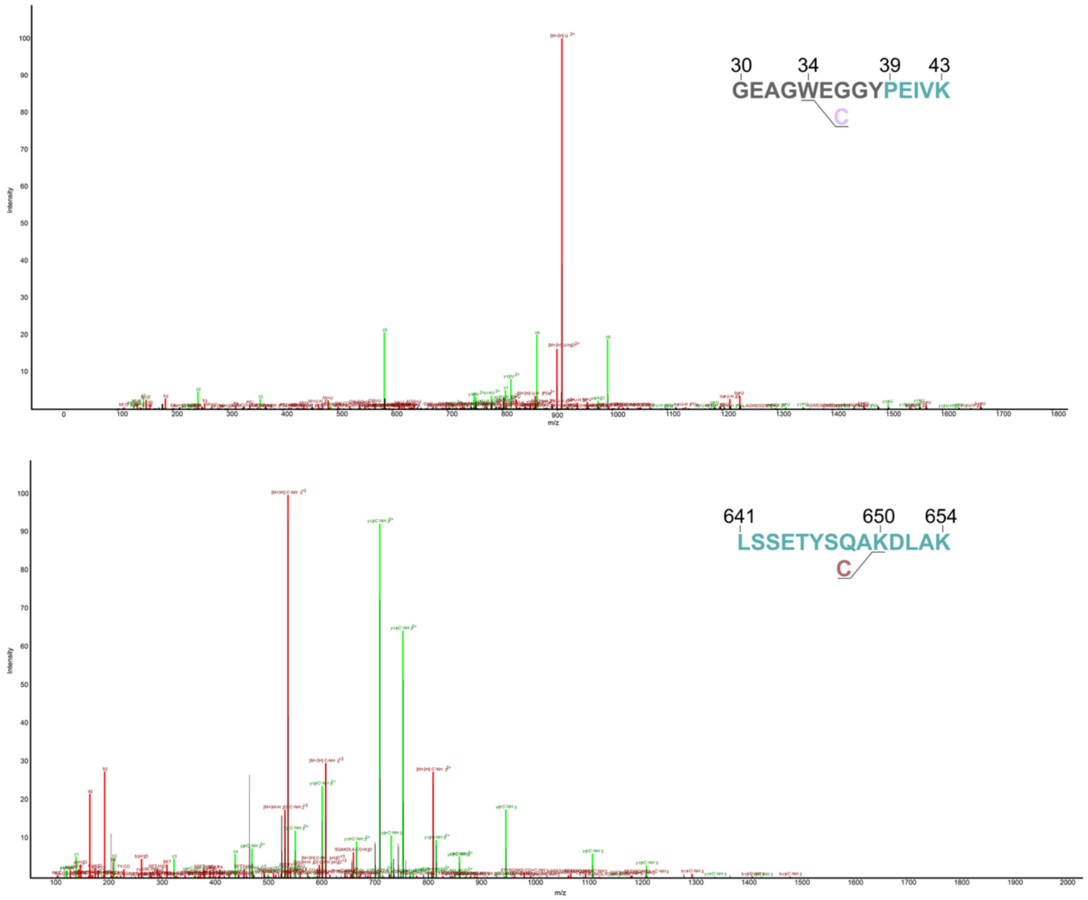

**Supplementary Data Fig.4 | Examples of protein-RNA UV crosslinking MS spectra.**
On top, exemplary crosslink spectrum match between the NSUN2 peptide GEAGWEGGYPEIVK (amino acids 30-43) and the nucleotide U from the $tRNA^{Asp}_{GUC}$. Both the Trp34 and the Tyr38 were identified as NSUN2 crosslink sites. On the bottom, exemplary crosslink spectrum match between the NSUN2 peptide LSSETYSQAKDLAK (amino acids 641-654) and the nucleotide C from the $tRNA^{Asp}_{GUC}$. Here, both the Tyr646 and the Lys650 were identified as NSUN2 crosslink sites. Precursor ions and their relative neutral losses (e.g., -H2O, -NH3, -H3PO4, -CO), as well as RNA bases/RNA bases+ribose reporter ions and b ions are depicted in red; y ions are displayed in green.

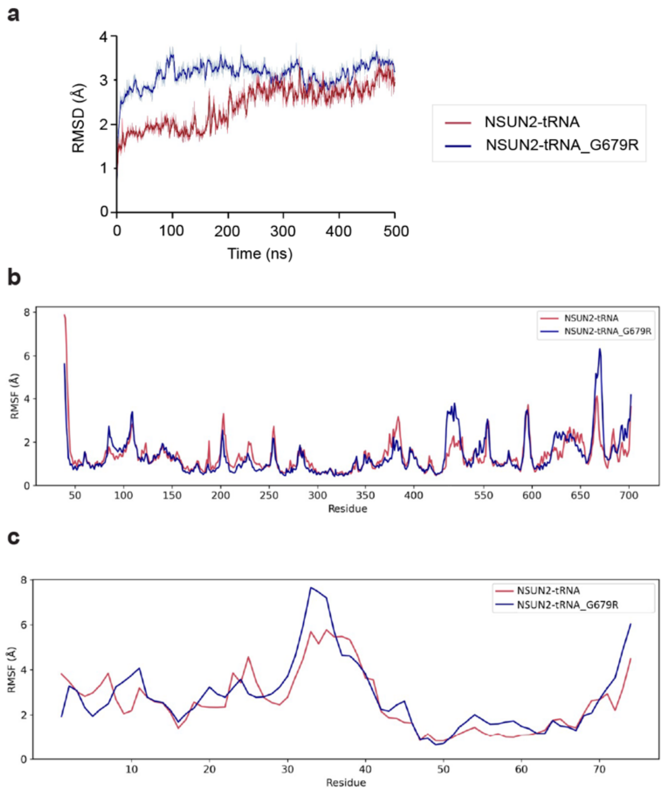

**Supplementary Data Fig.5 | MD simulations results of NSUN2 G679R mutation.**

**a,** Root-mean-square deviation (RMSD) of the NSUN2-tRNA complex (brick red) and the NSUN2(G679R)-tRNA mutant complex (dark blue) indicates that both systems remain structurally stable throughout the 500 ns molecular dynamics simulation. However, the mutant exhibits moderately higher RMSD values, suggesting reduced conformational stability compared to the non-crosslinked complex.

**b,** Superimposed per-residue backbone RMSF of the amino acid residues in the NSUN2-tRNA complex and its G679R mutant demonstrate consistently low average fluctuation values in both systems, indicating overall structural stability throughout the 500 ns molecular dynamics simulation. Higher fluctuations are observed only at a few terminal residues, whereas the core protein backbone remains stable over the course of the simulation. The plot further identifies localized regions exhibiting comparatively higher atomic mobility in both models.

**c,** Superimposed per-residue root mean square fluctuations (RMSF) of nucleotide residues in the NSUN2-tRNA complex and the corresponding G679R mutant complex obtained from molecular dynamics simulations. Residues 30-40 display higher flexibility, consistent with their localization within a loop region. The G679R substitution leads to increased fluctuations, indicating that the mutation enhances local structural dynamics of the NSUN2-tRNA assembly.

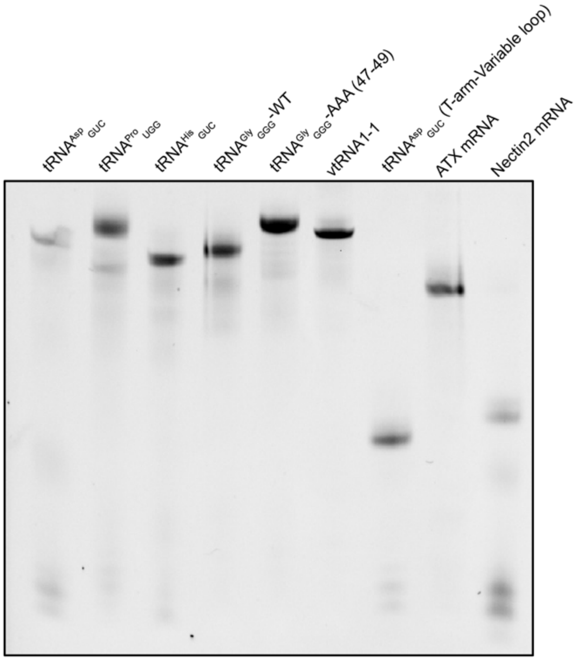

**Supplementary Data Fig.6 | RNA probes quality.**

Denaturing urea-PAGE of RNA probes obtained by *in vitro* transcription and used in EMSAs and covalent adduct assays.
